# A Vertebrate Toxin-Antidote System That Sabotages Mouse Embryogenesis

**DOI:** 10.64898/2026.03.04.709565

**Authors:** Duilio M.Z.A. Silva, Morgan Skinner, Takaya Totsuka, Takashi Akera

## Abstract

Toxin-antidote (TA) systems are selfish genetic elements that bias their own inheritance by coupling a toxin that kills offspring with an antidote that specifically rescues carriers. Although widespread across bacteria, archaea, fungi, plants, and invertebrates, TA systems have not been described in vertebrates. Here we report the first vertebrate TA system, which sabotages mammalian embryogenesis. We define HEX (Homogenously staining region-mediated Embryo eXecution) as a selfish element that biases its transmission through the female mouse germline. Crosses between HEX heterozygous females and wild-type males result in selective lethality of wild-type embryos, yielding preferential survival of HEX-bearing progeny. Using mouse genetics, embryo transfer, and zygote micromanipulation, we show that HEX operates through a canonical TA mechanism: the maternally deposited toxin SP100 induces genotoxic stress in embryos, while the linked antidote SP110 selectively rescues HEX-positive embryos. Both components are core factors of the interferon signaling pathway, revealing that HEX co-opts innate immune machinery to drive transmission bias. These findings establish a vertebrate TA system and demonstrate that selfish elements can repurpose fundamental cellular pathways to violate Mendelian inheritance, with profound consequences for female fertility.

## Introduction

Mendel’s Law of Segregation states that each parental allele is randomly transmitted to the next generation. Selfish genetic elements violate this Law by transmitting at a rate above 50%, often at a cost of organismal fitness, including reduced fertility. Although biased transmission of selfish elements has been described across diverse taxa, the underlying molecular mechanisms remain elusive in most systems.^1–6^ One strategy used by selfish elements is the toxin-antidote (TA) system, typically composed of a toxin and a genetically linked antidote.^7,8^ Selfish DNA encodes a toxin that is distributed to all the gametes in sexually reproducing organisms or to daughter cells in bacteria and archaea. The deposited toxin impairs the development of gametes, embryos, or cells that lack the selfish element, while the ones carrying the selfish DNA survive by expressing an antidote from the TA locus. This mechanism guarantees the increased transmission of the selfish element. TA systems were initially documented four decades ago in bacteria,^9,10^ and many have been found in archaea, fungi, invertebrate animals, and plants.^11–20^ However, whether such systems operate in vertebrates has remained unknown.

A Homogeneously Staining Region (HSR) on mouse chromosome 1 is a selfish element that exhibits biased transmission, specifically through the female germline.^21,22^ The HSR locus contains repetitive DNA composed of segmental duplications of at least two genes, *Sp100-rs* and *Sp110*, that are flanked by single-copy genes, *Sp100* and *Csprs,* on each end (Figure 1A).^23^ *Sp100-rs* is a chimeric gene that arose by fusion of the 5’ part of *Sp100* and *Csprs*.^23^ Speckled Protein (SP) family (e.g., SP100 and SP110) are chromatin readers that perform essential functions in diverse biological processes, including innate immunity,^24–28^ but their involvement in genetic conflict has not been explored. Whereas wild-type allele contains ∼50 copies of these sequences, the selfish allele expands at least to 800 copies, spanning over 100 Mb (Figure 1A).^29^ When HSR heterozygous (H/+) females are crossed with wild-type (+/+) males, approximately half of the +/+ embryos are lost, resulting in a strong transmission bias favoring H/+ offspring. Because HSR is a general cytogenetic term for chromosomal loci with homogeneous staining, we call this selfish element HEX (HSR-mediated Embryo eXecution), evoking a “hex”—a curse—that selectively eliminates wild-type embryos. HEX contrasts with typical selfish elements in the female germline that bias their own segregation to the egg,^30–39^ implying that HEX uses a previously unrecognized genetic cheating strategy in mammals.

**Figure 1.**
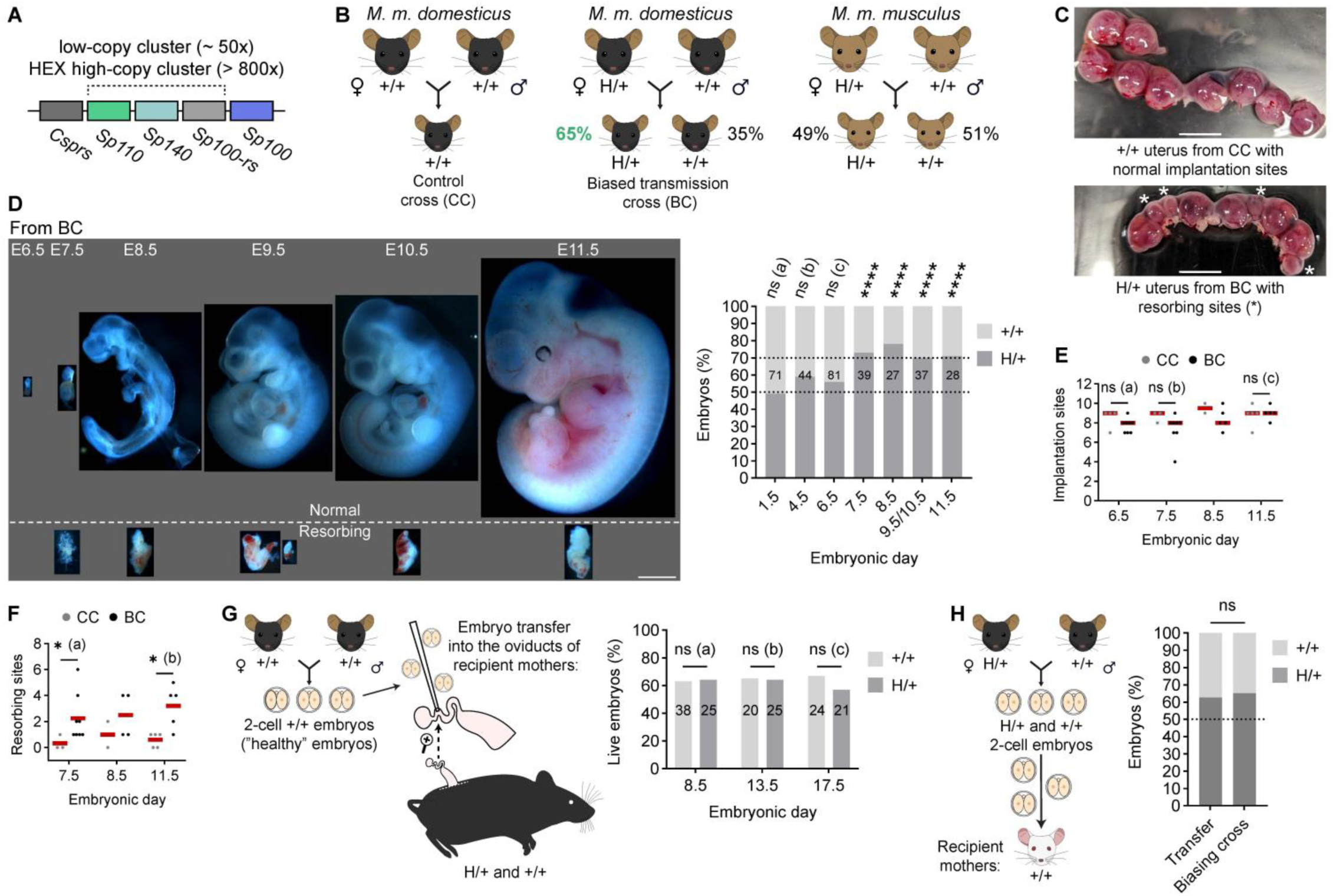
Selfish HEX kills wild-type embryos by causing internal issues. **(A)** Simplified diagram of the HEX locus (high-copy repeat) and wild-type low-copy repeat based on Weichenhan, D. et al.^56^ **(B)** Schematic of different crosses to study HEX transmission. **(C)** Uterus from the control cross (CC) with regular implantation sites and from the biasing cross (BC) with resorbing sites (*); scale bars, 500 *μ*m. **(D)** Time-course embryo dissection showing normal and resorbing embryos from the biasing cross; E, embryonic day; scale bar, 1 mm. Graph is a quantification of genotype ratios (H/+ versus +/+) at each time point; total number of embryos analyzed are indicated on top of each bar; Binomial test (two-sided) was used to examine deviation from the Mendelian expectation (50:50); ns (a) = 0.9204, ns (b) =0.0886, ns (c) = 0.2713, \*\*\*\**P* < 0.0001. **(E)** Numbers of implantation sites in the control cross (CC) and the biasing cross (BC) were quantified; n = 4, 8, 3, 8, 2, 5, 5, and 5 females from left to right groups; Mann-Whitney test (two-sided) was used for statistical analysis; ns (a) = 0.2788, ns (b) = 0.1636, ns (c) = 0.9524. **(F)** Numbers of resorbing sites in CC and BC were quantified; n = 3, 8, 2, 4, 5, and 5 females for E6.5 CC, E6.5 BC, E7.5 CC, E7.5 BC, E8.5 CC, E8.5 BC, E11.5 CC, and E11.5 BC, respectively; Mann-Whitney test (two-sided) was used for statistical analysis; *(a) *P* = 0.0485, *(b) *P* = 0.0317. **(G)** Schematic of the first embryo transfer experiment using healthy two-cell embryos from CC (left); percentages of live embryos relative to total implantation sites were quantified for each recipient mother’s genotype, +/+ and H/+, at different timepoints; number of embryos analyzed is indicated on top of each bar; Chi-square test (two-sided) was used for statistical analysis; ns (a) = 0.9458, ns (b) = 0.9445, ns (c) = 0.5109. **(H)** Schematic of the second embryo transfer experiment using two-cell embryos from BC (left); The genotype ratio of transferred embryos was compared to natural biasing cross (the data for natural cross is from Table 1); n = 51, 287 for Transfer and Biasing cross, respectively; Binomial test (two-sided) was used for statistical analysis; ns = 0.7692.

**Table 1.** HEX transmission bias in *Mus musculus domesticus* and *Mus musculus musculus* backgrounds. *Binomial test (two-tailed) for H/+ to +/+ ratios compared to Mendelian expectations (50:50).

| subspecies | dam genotype | sire genotype | H/+ progeny | +/+ progeny | total progeny | % H/+ | *P* value |
| --- | --- | --- | --- | --- | --- | --- | --- |
| *M. m. domesticus* | H/+ | +/+ | 187 | 100 | 287 | 65.2 | <0.0001* |
| *M. m. musculus* | H/+ | +/+ | 108 | 113 | 221 | 48.9 | 0.7373* |

## Results

### HEX drives post-implantation lethality of wild-type embryos

Previous study reported biased transmission of HEX, where chromosome 1 with HEX was part of the Robertsonian fusion with chromosome 18,^21^ raising the possibility that chromosomal rearrangement contributes to the phenotype. It is essential to disentangle their contribution from that of HEX, since Robertsonian fusions themselves can drive biased transmission in mice and humans.^30,40,41^ We therefore directly quantified transmission of the HEX locus in a standard *Mus musculus domesticus* (hereafter, *dom*) genetic background. Crosses between H/+ females and +/+ males (hereafter, biasing cross or BC in the Figures) yielded 65% H/+ offspring, significantly deviating from the expected Mendelian ratio (Figure 1B and Table 1). This biased transmission was independent of offspring sex (Table 2). These results confirm that selfish HEX autonomously biases its transmission, independent of Robertsonian fusion.

**Table 2.** HEX transmission bias in females and males of *Mus musculus domesticus*. *Chi-square test (two-tailed) for H/+ to +/+ ratios in females vs males.

| subspecies | dam genotype | sire genotype | H/+ females | +/+ females | total females | H/+ males | +/+ males | total males | *P* value |
| --- | --- | --- | --- | --- | --- | --- | --- | --- | --- |
| *M. m. domesticus* | H/+ | +/+ | 103<br>(70.1%) | 44 | 147 | 84<br>(60%) | 56 | 140 | 0.0736* |

Previous characterization found that +/+ embryos from the biasing cross die between day E4.5 (blastocyst) and E11.5.^21^ To precisely define the developmental window of lethality, we performed a high-resolution time course from E1.5 (two-cell) to E11.5 (Figures 1C and 1D). The time course embryo dissection revealed that E7.5 is the time when dead embryos at resorbing sites in the uterus are first observed (Figures 1D–1F). Notably, genotyping by qPCR (quantitative Polymerase Chain Reaction) and FISH (Fluorescence *in situ* Hybridization) showed that genotype ratios become skewed at this stage (Figure 1D, graph), suggesting that selective loss of +/+ embryos at E7.5 underlies HEX transmission bias. Since mouse embryos implant at E4.5–E5.5, selfish HEX kills genetically normal embryos shortly after implantation.

### Embryo lethality is intrinsic and independent of the uterine environment

Embryos could die either due to internal issues or due to harmful interactions with the maternal environment (i.e., external issues). Because the uterus and the dying embryos have distinct genotypes (H/+ and +/+, respectively), we wished to determine if genetic incompatibility upon implantation was responsible. We performed two distinct embryo transfer experiments. To test if the H/+ uterus can kill healthy ++ embryos, we transferred +/+ embryos that were never exposed to selfish HEX (from the +/+ x +/+ cross) to either H/+ recipient mothers or ++ control (Figure 1G). Transplanted embryos showed indistinguishable survival rates between H/+ and +/+ recipient mothers at E8.5, E11.5, and E17.5 (Figure 1G), implying that the H/+ uterus *per se* is not toxic to +/+ embryos.

To test if biased transmission is independent of uterus genotype, we transferred a mixture of H/+ and +/+ embryos from the biasing cross to +/+ uteri (Figure 1H). Embryo genotyping revealed a biased genotype ratio with 63% H/+ embryos, similar to the natural biasing cross (Figure 1H, graph). Furthermore, 66.5% of the implantation sites had dead embryos, consistent with the idea that embryo death is driving the transmission bias. Together, these results establish that HEX-mediated lethality is embryo-autonomous, indicating that the causal mechanism operates within the embryo rather than through maternal effects.

### HEX gene overexpression correlates with transmission bias

To date, the only established mechanism to sabotage/kill genetically wild-type cells/embryos by internal issues is the toxin-antidote (TA) system. The HEX locus encodes several genes (*Sp100*, *Sp110*, *Sp100-rs*, and *Csprs*) (Figure 2A) that could serve as a toxin or an antidote. In addition, the telomere-to-telomere mouse genome assembly^42^ found that another SP family gene, *Sp140*, is part of HEX repeats, and our qPCR analysis confirmed that *Sp140* is indeed amplified in mice with HEX (Figure S1). To test if these HEX genes contribute to biased transmission, we first analyzed their expression patterns by RT-qPCR (Reverse Transcription qPCR). RT-qPCR using liver tissues showed that *Sp100*, *Sp110*, and *Sp100-rs* genes were highly expressed in H/+ compared to +/+ (Figure 2B). Interestingly, *Sp100* shows increased expression in H/+ even though it is a single-copy gene in both genotypes. Similarly, we observed overexpression of HEX genes in H/+ oocytes compared to +/+ (Figure 2C). The results raise a possibility that the overexpression of HEX genes mediates biased transmission by embryo killing.

**Figure 2.**
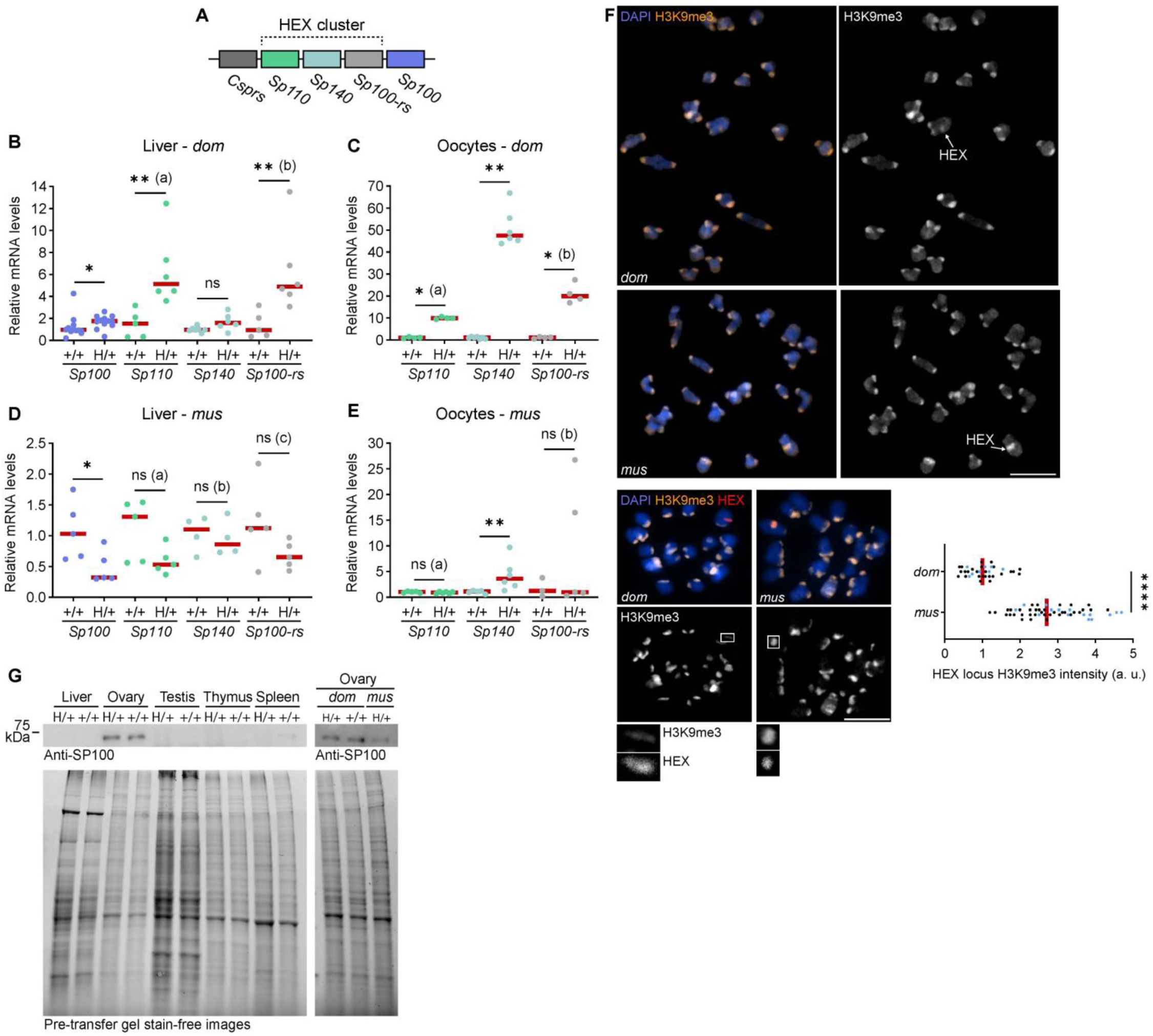
No HEX gene expression, no biased transmission. **(A)** Simplified diagram of HEX repeats based on Weichenhan, D. et al.^56^ **(B)** Transcript levels of HEX genes in +/+ and H/+ *Mus musculus domesticus* (*dom*) liver were analyzed by RT-qPCR; Mann-Whitney test (two-sided) was used for statistical analysis; n = 12, 12, 5, 6, 7, 6, 5, and 6 for samples from left to right groups. Data for *Sp100* was pooled from two independent experiments. \**P* = 0.0242, **(a) *P* = 0.0043, ns = 0.0513, **(b) *P* = 0.0087; red line, median. **(C)** Transcript levels of HEX genes in +/+ and H/+ *dom* oocytes analyzed by RT-qPCR; Mann-Whitney test (two-sided) was used for statistical analysis; n = 4, 4, 7, 6, 4, and 4 for samples from left to right groups. *(a) *P* = 0.0286, \*\**P* = 0.0012, *(b) *P* = 0.0286; red line, median. **(D)** Transcript levels of HEX genes in +/+ and H/+ *Mus musculus musculus* (*mus*) liver analyzed by RT-qPCR; Mann-Whitney test (two-sided) was used for statistical analysis; n = 5, 5, 5, 5, 4, 4, 5, and 5 for samples from left to right groups. \**P* = 0.0317, ns (a) *P* = 0.0952, ns (b) *P* = 0.8857, ns (c) *P* = 0.1508; red line, median. **(E)** Transcript levels of HEX genes in +/+ and H/+ *mus* oocytes analyzed by RT-qPCR; Mann-Whitney test (two-sided) was used for statistical analysis; n = 5, 6, 5, 6, 3, and 5 for samples from left to right groups. ns (a) = 0.329, \*\**P* = 0.0087, ns (b) = 0.7857; red line, median. In BE, comparisons are made within each gene, and the values are not intended for comparisons across different genes. **(F)** H/+ oocytes from *dom* and *mus* were fixed at prometaphase I and stained for H3K9me3 (top). H3K9me3 signal intensities at the HEX locus were quantified; datapoints from two independent experiments are indicated by dots with distinct colors; n = 34 and 51 for *dom* and *mus*, respectively; Welch’s t test (two-sided) was used for statistical analysis; \*\*\*\**P* < 0.0001; red line, median. Images are sum intensity Z-projections showing all chromosomes; scale bar, 10 *μ*m. H/+ oocytes from *dom* and *mus* were fixed at prometaphase I and stained for H3K9me3, followed by FISH using the HEX probe (bottom). Images are sum intensity Z-projections showing all chromosomes; scale bar, 10 *μ*m, scale bar of enlarged insets, 1 *μ*m. **(G)** Western blot of SP100 using tissue samples from H/+ and +/+ *dom* (left) and *dom* and *mus* (right). As a loading control, stain-free gel images are shown; see Figure S2 for gel source data.

Some meiotic drive systems in *Mus musculus domesticus* become suppressed when introduced to another subspecies, *Mus musculus musculus* (hereafter, *mus*).^35,43^ We tested if this applies to our system by backcrossing the HEX locus into the *mus* background for six generations and found that the HEX cheating was completely suppressed (Table 1, *M. m. musculus*). RT-qPCR analyses using *mus* liver and oocytes showed that HEX genes do not overexpress in H/+ compared to +/+ (Figures 2D and 2E), consistent with the idea that HEX gene overexpression contributes to biased transmission.

The HEX locus shows a distinct C-band staining pattern,^44^ indicative of heterochromatin.^45^ This raised the possibility that heterochromatin regulation modulates HEX gene expression profiles in *dom* vs *mus*. We examined H/+ oocytes from *dom* and *mus* by immunostaining and found that levels of histone H3K9me3, a heterochromatin marker, are significantly higher at the HEX locus in *mus* compared to *dom*, consistent with the HEX gene silencing in the *mus* background (Figure 2F). These results implicate epigenetic silencing as a natural defense against selfish elements.

### Toxin candidate is equally distributed to all embryos

Embryo transfer experiments and gene expression analyses imply that HEX genes regulate biased transmission, possibly through a TA system. There are at least three main requirements for a TA system to operate in the female germline: (1) the toxin needs to be produced during oogenesis and be deposited to all embryos, (2) the antidote needs to function after fertilization, specifically in the embryos with the selfish element, and (3) when the toxin alone is present in embryos, it must sabotage embryogenesis. It is critical to examine which HEX genes satisfy these requirements to operate a TA system during mammalian reproduction. A key clue emerged from studies of innate immunity in human cells, where SP100 overexpression triggers DNA damage and cell death within the interferon signaling cascade.^28^ Notably, co-expression of SP110 suppresses this cytotoxicity.^28^ These observations led us to propose that HEX co-opts innate immune factors in the female germline to challenge Mendelian inheritance, with SP100 acting as a toxin and SP110 as its antidote.

To test the toxin (SP100)–antidote (SP110) hypothesis for HEX cheating, we first examined SP100 protein levels across different tissues. We found that the major isoform of the SP100 protein (68 kDa) is highly expressed in ovaries, compared to liver, spleen, thymus, and testis (Figures 2G and S2). This result is consistent with the first requirement for a TA system, that the toxin is produced during oogenesis. The western blot analysis also showed lower SP100 levels in H/+ *mus* ovaries compared to *dom* (Figure 2G, right), consistent with the idea that the *mus* background suppresses the expression of HEX genes to cancel biased transmission. H/+ and +/+ ovaries showed similar SP100 protein levels, but ovaries are composed mostly of non-gamete cells. Therefore, we decided to analyze SP100 levels in more detail at a cellular level, focusing on oocytes and embryos (see below).

To directly visualize how the toxin candidate, SP100, is deposited in embryos, we analyzed SP100 levels and its localization pattern in oocytes, eggs, and two-cell embryos. Interestingly, meiosis I oocytes from H/+ females did not show higher SP100 levels compared to +/+, but meiosis II eggs and two-cell embryos from H/+ females (biasing cross, BC) showed increased SP100 levels compared to those from +/+ (control cross, CC) (Figures 3A and 3B). Crucially, SP100 levels were equally high in the embryos from the biasing cross regardless of embryo genotype (H/+ and +/+) (Figure S3), consistent with the hypothesis that SP100 toxin is uniformly distributed to all eggs and embryos when cheating happens.

**Figure 3.**
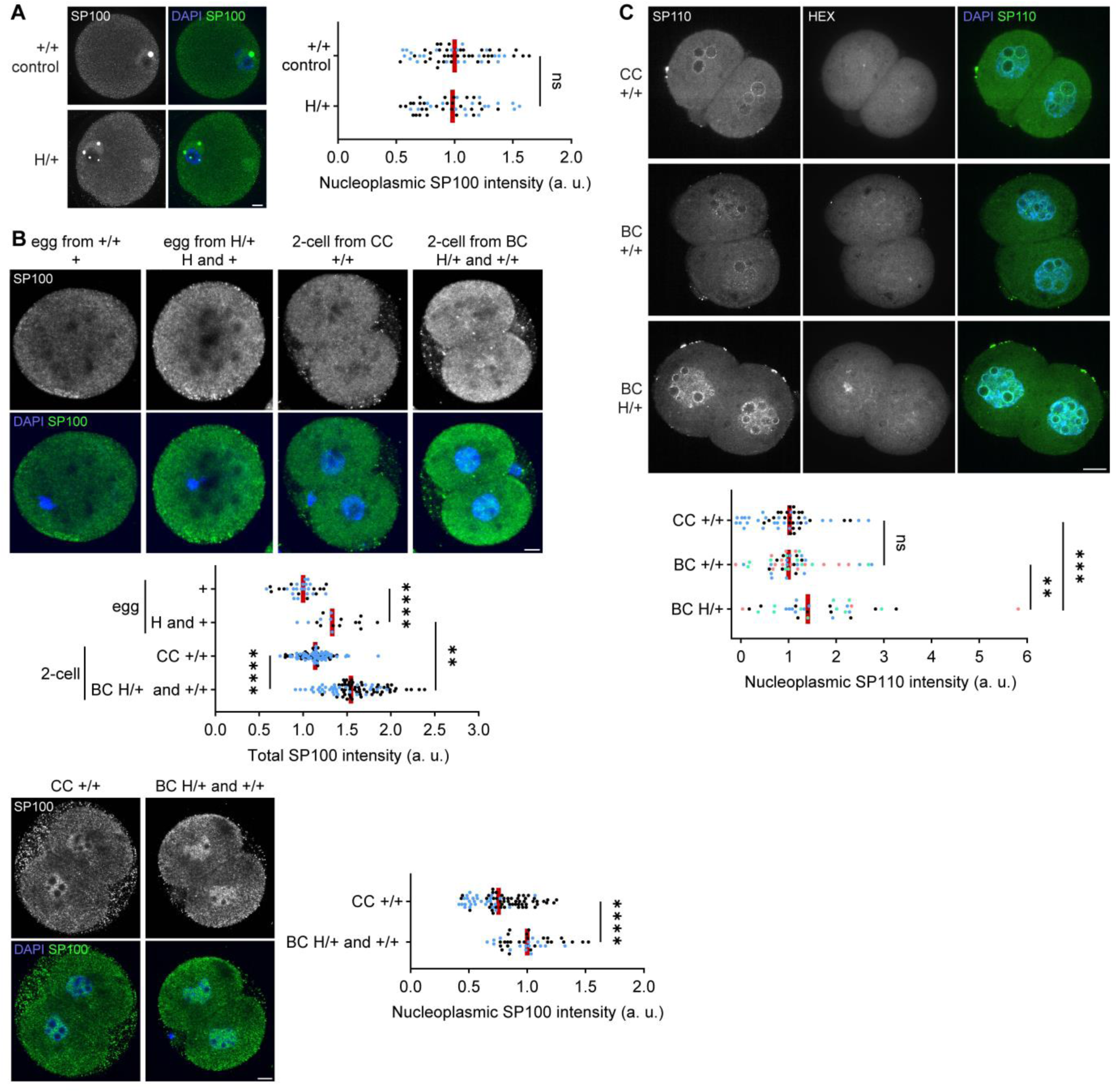
Toxin is deposited in all embryos while the antidote is private to HEX embryos. **(A)** Oocytes were fixed at the germinal vesicle (GV) stage and stained for SP100. The signal intensities were quantified; n = 58 and 47 for +/+ control and H/+, respectively; Mann-Whitney test (two-sided) was used for statistical analysis; ns = 0.3912; red line, median; dots, individual cells from two-cell embryos. Images are maximum intensity Z-projections; scale bar, 10 *μ*m. **(B)** metaphase II eggs and two-cell embryos were fixed and stained for SP100. The total signal intensities in each cell were quantified (top) (n = 31, 17, 100, and 92 for + egg, H and + egg, CC +/+ 2-cell, BC H/+ and +/+ 2-cell, respectively). Welch’s t test (two-sided) was used for statistical analysis; \*\*\*\**P* < 0.0001, \*\**P* = 0.0063; red line, median; dots, individual cells. Nucleoplasmic signal intensities were quantified in two-cell embryos (bottom) (n = 80 and 42 for CC +/+ and BC H/+ and +/+, respectively); Mann-Whitney test (two-sided) was used for statistical analysis; \*\*\*\**P* < 0.0001; red line, median; dots, individual cells. Images are maximum intensity Z-projections; scale bar, 10 *μ*m. **(C)** Embryos were fixed at the late two-cell stage, stained for SP110, and genotyped by FISH with the HEX probe. The signal intensities were quantified (n = 50, 47, and 33 for CC +/+, BC +/+, and BC H/+, respectively). Mann-Whitney test (two-sided) was used for statistical analysis; ns = 0.3368, \*\**P* = 0.0037, \*\*\**P* = 0.0004; red line, median; dots, individual cells. This experiment was independently repeated two times for CC +/+ and four times for BC +/+ and BC H/+. Images are maximum intensity Z-projections; scale bar, 10 *μ*m. Data points from two or more independent experiments are indicated by dots with distinct colors.

### Antidote candidate is specifically upregulated in embryos with HEX

A second requirement for TA systems is that the antidote needs to function after fertilization, specifically in the embryos with the selfish element. Indeed, the antidote candidate, SP110, is an established two-cell embryo marker, responding to the zygotic genome activation (ZGA).^25^ This expression timing is ideal for SP110 to interact with the maternally deposited toxin candidate, SP100, to suppress its toxicity. To test if embryos with selfish HEX show higher SP110 levels compared to the dying +/+ embryos, we analyzed late two-cell embryos that have likely completed ZGA. Our analysis showed that H/+ embryos have significantly higher nuclear SP110 levels compared to +/+ embryos in the biasing cross (Figure 3C), consistent with the hypothesis. The SP110 level in +/+ embryos from the biasing cross was similar to that of +/+ embryos from the control cross (Figure 3C), suggesting that while +/+ embryos show basal SP110 expression, H/+ embryos overexpress SP110, likely due to their higher gene-copy number, and thus specifically neutralize the toxic effect. These results show that SP110 meets the requirement to serve as an antidote in the HEX system.

Our model predicts that HEX-mediated toxicity is suppressed upon paternal transmission. While the SP100 toxin is maternally deposited into all embryos, the antidote SP110 is expressed after fertilization. Thus, embryos inheriting HEX—even from sperm—can express SP110 and neutralize the toxin. Consistent with the model, previous study have shown that transmission bias is canceled when crossing H/+ females with H/H males.^21^ Furthermore, the asymmetric contribution of toxin explains why transmission bias is restricted to the female germline.

### Dying +/+ embryos have increased DNA damage

A third requirement for a TA system is that a toxic effect occur in embryos when the toxin is not suppressed by an antidote. Since SP100 overexpression causes increased DNA damage in human cells,^28^ we asked if dying +/+ embryos exhibit elevated DNA damage. We examined a DNA damage marker, 53BP1,^46^ in embryonic cells from day E6.5, one day prior to the onset of +/+ embryo death. The immunofluorescence analysis showed that +/+ embryos from the biasing cross have higher levels of DNA damage compared to H/+ embryos from the same cross and +/+ embryos from the control cross (Figure 4A), linking SP100 activity with genotoxicity and embryo killing.

**Figure 4.**
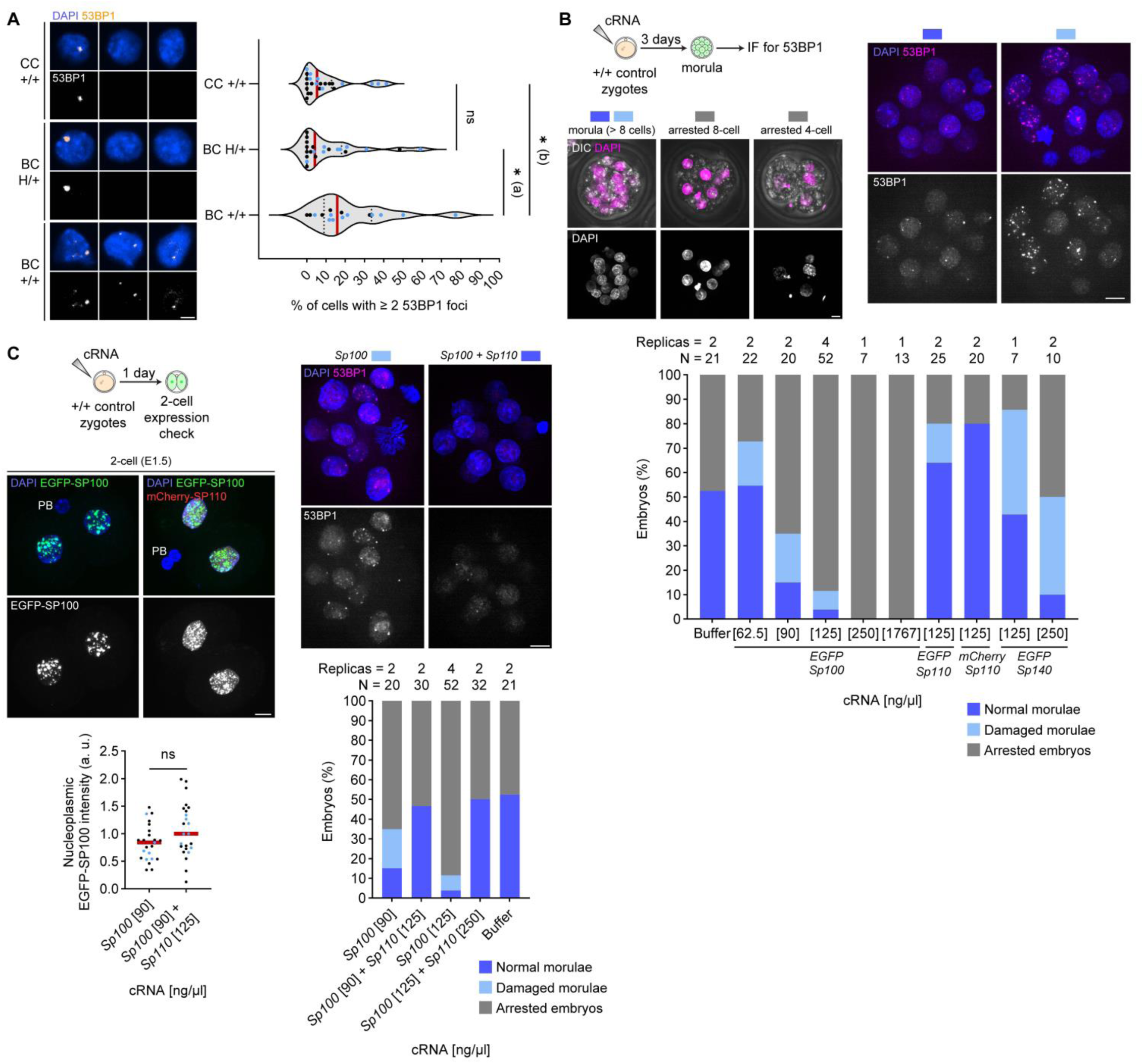
SP100 and SP110 consist a minimal toxin-antidote module, sabotaging mammalian embryogenesis. **(A)** Embryos from control cross (CC) and biasing cross (BC) were fixed at E6.5 and stained for 53BP1. Percentages of cells with more than two 53BP1 foci in each embryo were quantified; Mann-Whitney test (two-sided) was used for statistical analysis; ns = 0.9691, *(a) *P* = 0.0254, *(b) *P* = 0.0102; red line, median; dots, percentage per embryo. Data points from two independent experiments are indicated by dots with distinct colors (n = 24, 22, 16 embryos for CC +/+, BC H/+, and BC +/+, respectively). **(B)** Zygotes were microinjected with cRNA, cultured for three days (E3.5), and fixed to stain for 53BP1. Embryos at the morula stage (> eight cells) were classified as normal morulae (< two cells within the embryo with five or more 53BP1 foci, dark blue) or damaged morulae (two or more cells with five or more 53BP1 foci, light blue), and embryos with eight cells or fewer were classified as arrested embryos (grey). Proportions of embryos in each category were quantified; sample size and the number of independent experiments are shown above each bar in the graph. **(C)** Embryos expressing EGFP-SP100 and mCherry-SP110 were fixed at the two-cell stage and counterstained with DAPI. EGFP-SP100 nucleoplasmic signal intensities were quantified as a readout of expression levels; datapoints from two independent experiments are indicated by dots with distinct colors; n = 23 and 24 for *Sp100* [90] and *Sp100* [90] + *Sp110* [125], respectively; Welch’s t test (two-sided) was used for statistical analysis; ns = 0.0625; red line, median; dots, individual cells. Similarly, embryos expressing EGFP-SP100 and mCherry-SP110 were cultured for three days, fixed and stained for 53BP1, and categorized as in B (bottom); note that data for *Sp100* [90], *Sp100* [125], and Buffer are from the graph in B; sample size and the number of independent experiments are shown above each bar. Images in this figure are maximum intensity Z-projections except for the ones with DIC, which are standard deviation intensity Z-projections; PB, polar body; scale bar, 10 *μ*m.

### SP100 and SP110 compose a minimal TA module

To directly test if SP100 and SP110 are sufficient to operate a TA system during mouse embryogenesis, we ectopically expressed them by microinjecting their cRNA into +/+ zygotes from the control cross and cultured the embryos for three days (E3.5, the morula stage) (Figure 4B). SP100 alone potently disrupted embryogenesis in a dose-dependent manner, inducing developmental arrest and DNA damage. Co-expression of SP110 suppressed these defects without altering SP100 levels, demonstrating functional neutralization (Figures 4B–4C). We noticed that SP100-expressing embryos often show DNA bridges, which may be the origin of the DNA damage (Figure S4), consistent with the observation in human cells.^28^ These results establish SP100 and SP110 as a minimal toxin–antidote pair capable of recapitulating HEX activity.

We also tested whether other genes on the HEX locus, *Sp100-rs* and *Sp140*, impact embryogenesis when overexpressed. While SP100-rs overexpression did not impact embryogenesis (Figure S5A), SP140 overexpression had mild toxic effects (Figure 4B), possibly contributing as a secondary toxin. However, SP140 levels are similar between the control and biasing crosses in two-cell embryos (Figure S6A), suggesting that SP140 is not a primary toxin and that SP100–SP110 constitutes the core functional module.

## Discussion

### A vertebrate toxin-antidote system that repurposes innate immune mechanisms

Our findings represent the first canonical toxin-antidote system in vertebrates and the first genetic cheating mechanism that exploits the DNA damage pathway. Together with a previous report investigating a TA system in *C. elegans* that delays larval development,^20^ subverting embryogenesis may be a common strategy for TA systems in animals. Notably, both toxin and antidote for the HEX system derive from the interferon signaling pathway, suggesting that innate immune components can be repurposed to enforce selfish transmission. This study expands the conceptual framework of genetic conflict by showing that core cellular defense pathways can be redirected toward intragenomic competition.

### Temporal gating of toxicity

Our HEX TA model (Figure 5) proposes that H/+ females poison their embryos by increasing SP100 protein levels in meiosis II eggs. Some mRNAs that are critical for oocyte maturation and early development exhibit dormancy in oocytes and only initiate translation in eggs and embryos.^47^ *Sp100* mRNA may be employing a similar mechanism to produce toxin at the right place at the right time. It is also important to consider why the SP100 toxin, which is already upregulated in the two-cell stage, kills wild-type embryos only at day E6.5–7.5. We speculate on two possibilities. First, SP100 may lack critical cofactors to induce DNA damage during early embryogenesis. Indeed, human SP100 requires a nuclear body, the PML body, to induce DNA damage,^28^ and the PML body is absent from oocytes and early embryos and only starts forming at the morula stage.^48^ This delayed formation of the PML body can explain why SP100 does not immediately kill embryos after fertilization. Second, SP100 may cause mild DNA damage in early embryos, which only triggers death after implantation. Consistent with this model, previous work has shown that embryos become particularly sensitive to mild DNA damage, resulting in apoptosis at E6.5 when gastrulation takes place.^49^ Furthermore, multiple mutant mouse lines with elevated DNA damage during embryogenesis show lethality at E6.5–E7.5,^49–53^ consistent with our model that HEX kills embryos via DNA damage.

**Figure 5.**
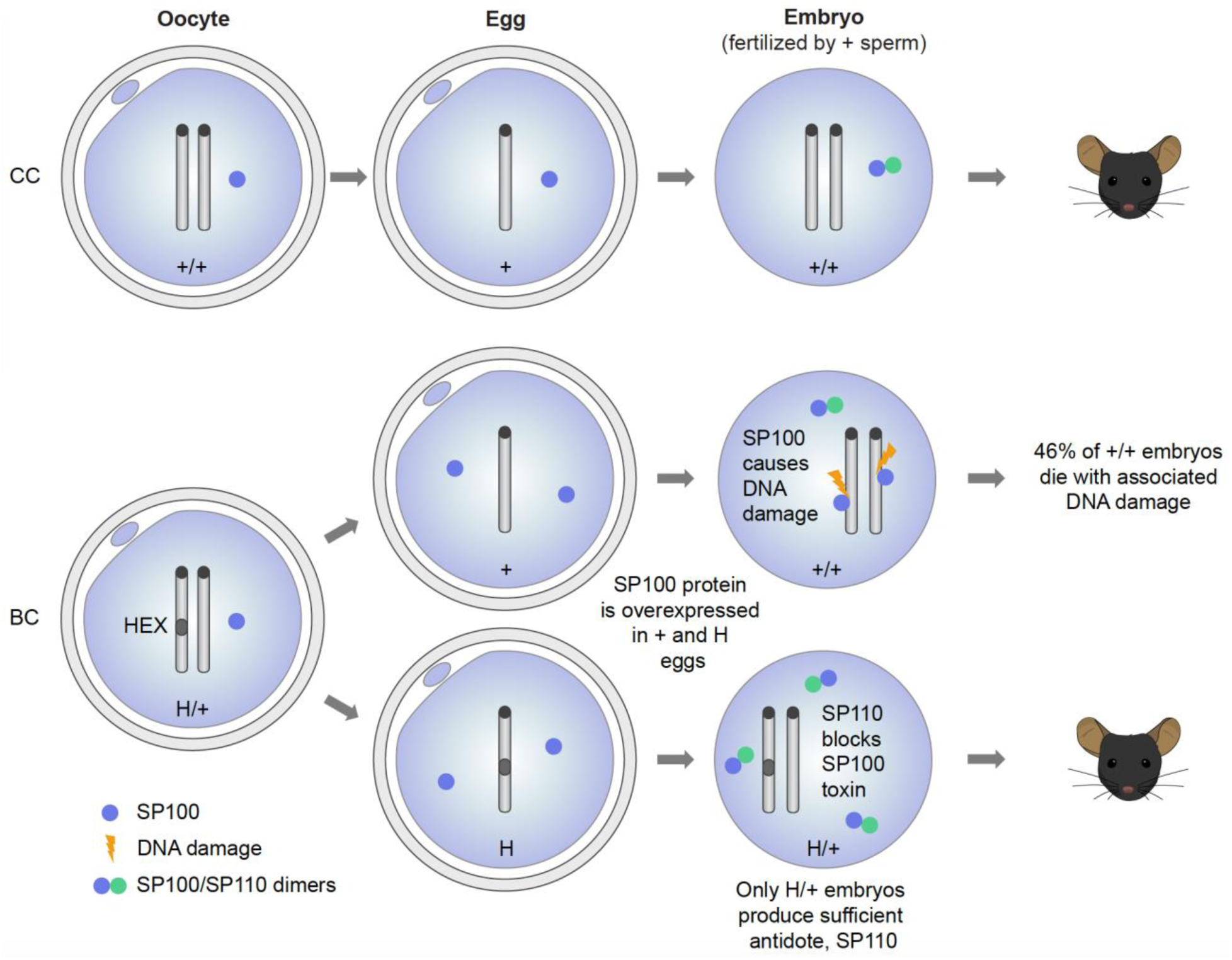
HEX represents the first canonical toxin-antidote system in vertebrates. SP100 toxin becomes enriched in eggs and early embryos from H/+ females. SP100 levels are elevated in all embryos from biasing crosses (BC), regardless of genotype. In contrast, SP110 antidote is selectively upregulated in H/+ embryos following fertilization. This differential expression provides a mechanism by which only HEX-positive embryos neutralize genotoxicity. Although the t-haplotype meiotic drive system in male mice has occasionally been categorized as a TA-like system, it involves multiple distorters and a responder, and therefore mechanistically, it is a distorter-responder system. In contrast, HEX employs a tightly linked two-gene toxin-antidote module, characteristic of a canonical TA system. In human cells, the SP100-SP110 interaction via the CARD domain is crucial for neutralizing the cytotoxicity of SP100 overexpression (Figure S6B).^28^ Neutralization through the toxin-antidote direct interaction is another feature of canonical TA systems. Therefore, we believe HEX represents the first canonical TA system in vertebrates.

### Evolutionary dynamics of suppression and innovation

HEX activity is fully suppressed in the *M. m. musculus* background likely through heterochromatin-mediated silencing, illustrating a host defense mechanism against selfish elements (Table 1 and Figure 2).^54^ Some *M. m. musculus* populations carry a structural variant of HEX element called “double-band”, composed of two conspicuous heterochromatic bands originating from a paracentric inversion involving the HEX locus. Remarkably, the double-band cheats during meiosis by biased segregation to the egg.^43,55^ These observations raise an intriguing possibility that HEX escaped the suppression in *M. m. musculus* by evolving a new cheating strategy in a distinct arena: asymmetric female meiosis. This highlights the evolutionary plasticity of genetic conflict systems.

TA systems cause intragenomic conflict between the selfish element and the rest of the genome, facilitating genome innovation. Thus, investigating the HEX TA system would also provide a unique window to dissect how such conflicts have shaped mammalian genome, development, and reproduction.

### Limitations of the Study

Although our RT-qPCR analyses distinguish *Sp100* and *Sp100-rs* transcripts, the SP100 antibody used in this study may also recognize SP100-rs, if it is expressed as a protein. Thus, the immunofluorescence signals in Figures 3A and 3B could, in principle, include contributions from SP100-rs. However, SP100-rs lacks the C-terminal region of SP100, including the nuclear localization signal (NLS) (Figure S6B), and correspondingly displays diffuse localization in embryos when expressed as a GFP fusion (Figure S5B). Therefore, we consider that the increased nuclear enrichment of SP100 in embryos from H/+ females is indeed the full-length SP100 with intact NLS rather than SP100-rs (Figure 3B).

## Resource availability

### Lead contact

Further information and requests for resources and reagents should be directed to and will be fulfilled by the lead contact, Takashi Akera.

### Materials availability

The microscopy and western blot data required to reproduce the results in the study will be publicly available in Figshare upon publication. Any additional information required to reanalyze the data reported in this paper is available from the corresponding authors upon request.

## Data and code availability

– All original codes are available in this study’s Supplementary Text.
– Any additional information required to reanalyze the data reported in this paper is available from the lead contact upon request.

## Acknowledgments

We thank M. Lichten and J.L. Mueller for comments on the manuscript, Akera lab members for discussion, N. Altan-Bonnet, Y. Mukoyama, N. Mizuno, J.R. Hogg, and K. Sadtler for access to their laboratory equipment, G. Arora, R. Pavani, F.E. Clark, L.F. Rosin, V. Sundaresan, and W. Li for technical advice, A. Nussenzweig for the 53BP1 antibody, R.E. Vance for the SP140 antibody, NHLBI Transgenic Core (C. Liu) for embryo transfer experiments, and NIH building 14F animal facility staff for the care with our mouse colony, especially the “jumpy” mice. The contributions of the NIH authors are considered Works of the United States Government. The findings and conclusions presented in this paper are those of the authors and do not necessarily reflect the views of the NIH or the U.S. Department of Health and Human Services.

## Author contributions

Conceptualization: DMZAS and TA; Methodology: DMZAS, MS, TT; Investigation: DMZAS, MS, TT, TA; Funding acquisition: TA; Supervision: TA; Writing – original draft: DMZAS; Writing – review & editing: DMZAS, MS, TT, TA.

## Experimental model details

### Mouse strains and crossing schemes

Mouse strains were purchased from the Jackson Laboratory (B6.Cg-Is(HSR;1)1Lub/J, stock# 001914, *Mus musculus domesticus* with the HEX locus on chromosome 1, C57BL/6J, stock# 000664, *Mus musculus domesticus*, PWD/PhJ, stock# 004660, *Mus musculus musculus*) or Charles River (CD-1^®^ IGS, stock# 022, *Mus musculus domesticus*). The HEX locus was introduced to the *M. m. musculus* background by backcrossing H/+ *M. m. domesticus* mice with +/+ *M. m. musculus* (PWD/PhJ) mice for six generations. HEX transmission was measured by crossing H/+ *dom* females and +/+ *dom* males or by crossing H/+ *mus* females from the backcrosses and +/+ *mus* males. Mice were maintained in an animal facility under a 12-hour light/dark cycle at room temperature, with minimal disturbance and humidity ranging from 30% to 70% depending on the season. All animal experiments were conducted in accordance with the National Institutes of Health guidelines and approved by the Animal Care and Use Committee (Animal Study Proposal Numbers: H-0327 and H-0125).

## Method details

### Mouse oocyte and egg collection and culture

For oocyte collection, 6-12 week old *M. m. domesticus* females were primed with 7.5 U of pregnant mare serum gonadotropin (PMSG, Medix Biochemica, cat# 493-10) 44-48 hour before collection, while *M. m. musculus* females were primed with 5 U of CARD HyperOva (Cosmo Bio, cat# KYD-010-06-EX) 44-48 hour before collection. For metaphase II egg collection, the females were also primed with 7.5 IU of hCG (Millipore, Cat# 9002–61-3 C1063) 48 hours after the PMSG injection and euthanized 17 hours after the hCG injection. Germinal vesicle (GV) oocytes were collected in M2 media (Sigma, cat# M7167) with 2.5 mM milrinone (Sigma, cat# 475840) to inhibit meiotic resumption. Oocytes were then transferred to M16 media (Millipore, cat# M7292) with 2.5 mM milrinone covered with parafin oil (Nacalai, cat# NC1506764) in a humidified atmosphere of 5% CO_2_ in air at 37°C. For prometaphase I analysis (Figure 2F), milrinone was washed out to allow meiotic resumption. The oocytes were then fixed 3 hours after milrinone washout. Eggs were collected in M2 media from the ampulla of 4-12 week old females, transferred to M16 media covered with paraffin oil, and incubated at 37°C in a humidified atmosphere of 5% CO_2_ in air. Oocyte and egg manipulations were performed in a Petri dish (Falcon, cat# 351007).

### *In vitro* fertilization to obtain zygotes

Zygotes were obtained by IVF following the Nakagata method,^57^ using +/+ *dom* sperm and H/+ or +/+ *dom* eggs, except that eggs were collected in HTF media (Millipore, cat# MR-070-D). Cumulus cells and sperm were washed out 4-6 hours after fertilization in HTF media, and zygotes were transferred to KSOM media (Millipore, cat# MR-106) under paraffin oil. Zygote pronuclei were examined with an M80 stereo microscope (Leica, cat# 01-672-670) with a Thermo Plate TPX (Leica). Embryos with anormal pronuclei numbers and unfertilized eggs were discarded. Zygotes were cultured at 37°C in a humidified atmosphere of 5% CO_2_ in air to reach various stages.

### Post-implantation embryo collection

Embryos were dissected from H/+ *dom* pregnant females naturally mated with +/+ *dom* males. Dissections were performed during embryonic days E6.5 to E11.5 following standard laboratory protocols.^57,58^ Embryos were placed in HBSS media (Gibco, cat# 14175095) and imaged using the MZ16F stereomicroscope (Leica) and the Leica Application Suite X software. After imaging, embryos were frozen at −70°C until genotyping (see below).

### DNA isolation and genotyping by qPCR

Genomic DNA (gDNA) was extracted from tail snips following the HotSHOT protocol.^59^ Samples were genotyped by measuring the relative quantification (RQ) of the *Sp100-rs* gene by qPCR. RQ was estimated through the 2^−Δ*C*^_T_ method^60^ using the *36b4* gene as a reference (Table S1). Primers were designed using a modified version of Primer3 2.3.7^61^ implemented in Geneious 10.2.6 (Biomatters) (Table S1). The presence of the *Sp140* gene in the HEX locus (Figure S1) was also tested with a specific primer set (Table S1) using the *36b4* gene as a reference. Quantitative PCR was performed on a LightCycler 96 (Roche). Reactions were performed at a final volume of 10 μl, with 60 ng of gDNA, 5 μl of iTaq Universal SYBR Green Supmix (Bio-Rad, cat# 1725121), and 1 μl of each 5 μM primer. Cycle conditions were 95°C for 10 min; 40 cycles of 95°C for 5 s and 60°C for 20 s. Target and reference genes were analyzed simultaneously in triplicate. The specificity of the PCR products was confirmed by dissociation curve analysis. We estimated the amplification efficiency (E) of each primer set by standard curve performed on a 5-fold dilution series of gDNA. Amplification efficiencies were E*Sp100-rs* = 100% and E*36b4* = 96%, which are acceptable to use for the 2^−Δ*C*^_T_ method without correction. We classified animals as wild-type, heterozygous for HEX, and homozygous for HEX, comparing their RQ values with those obtained from animals with known genotypes. Animals with inconsistent RQ values were excluded from any following experiments. E1.5 to E7.5 embryos were genotyped by FISH using an HEX probe (see below).

### Oligopaint design and genotyping by FISH

We manually designed a set of probes for the HEX locus. Probes were designed at the *Sp100-rs* and *Sp110* exons and were synthesized by Integrated DNA Technologies. E1.5-E4.5 embryos produced by natural crosses were spread onto microscope slides based on the standard protocol.^62,63^ Briefly, embryos were treated with a hypotonic solution (0.1% sodium citrate in 0.2 mg/ml BSA), transferred to a hypotonic solution drop onto the microscope slide, and air dried. Just before completely dried, 8 µl of spreading solution (0.1% Tween-20 in 0.01 N HCl) was added over the cells and allowed to dry for about 1 hour. Cells were then fixed using a 1:3 mixture of acetic acid/methanol and let air dry. The FISH process was performed following the procedures described in Clark et al^64^ using 100 pmol of the HEX primary probe. E6.5 and E7.5 embryos were spread and genotyped by FISH as described in “Immunostaining and FISH on E6.5 embryos” (see below).

### Western blot

Liver, ovary, testis, thymus, and spleen samples were dissected, flash frozen in liquid nitrogen, and stored at −70°C. To extract protein from dissected tissues, 50 mg of each sample were homogenized using RIPA buffer (150 mM NaCl (Millipore, cat# 116224), 0.5% Sodium deoxycholate (Biochemica, cat# A1531), and 50 mM Tris, pH 8.0 (KD Medical, cat# RGF-3360), and 1% Triton X-100) supplemented with the protease inhibitor cocktail (MedChem Express, cat# HY-K0010). Extracts were mixed with Laemmli buffer (BioRad, cat# 1610737) containing 2-mercaptoethanol (VWR Life Sciences, cat# 97064-590) and denatured at 95°C for 5 min. Samples were separated on 10% Mini-Protean TGX Stain-Free Protein Gels (Bio-Rad, cat# 4568035) to visualize the total proteins with the ChemiDoc MP Imaging System (Bio-Rad, cat# 12003154). Proteins were then transferred to the PVDF membrane (Bio-Rad, cat# 1704156) with the Trans-Blot Turbo System (Bio-Rad, cat# 1704150). The membrane was blocked for 1 hour at RT with 2% Membrane Blocking agent (Cytiva, cat# RPN2125), incubated overnight at 4°C with the mouse anti-human SP100 (1:1000, Santa Cruz, cat# sc-293458), and washed three times for 15 min each with the TBST buffer (20 mM Tris, pH 7.6, 150 mM NaCl, and 0.1% Tween-20). The membrane was then incubated for 1 hour with the Goat Anti-Mouse IgG-heavy and light chain cross-adsorbed Antibody HRP conjugated (1:10,000 Bethyl, cat# A90-516P), followed by three 15 min washes with the TBST buffer. SP100 protein was visualized using the ECL detection reagents (Bio-Rad, cat# 1705062) and ChemiDoc MP Imaging System.

### Immunostaining of whole oocytes, eggs, and two-cell embryos

Oocytes at the germinal vesicle stage (GV) or prometaphase I (3 hour from milrinone washout), meiosis II eggs, and early two-cell embryos were fixed in freshly prepared 2% paraformaldehyde (Electron Microscopy Sciences, cat# 15710) in 1x PBS (Quality Biological, cat# 119-069-101CS) with 0.1% Triton X-100 (Millipore, cat# TX1568–1) for 20 min at room temperature (RT), washed in blocking solution (0.3% BSA (Fisher bioreagents, cat# BP1600–100) and 0.01% Tween-20 (ThermoFisher, cat# J20605-AP) in 1x PBS) for 10 min at RT, permeabilized in 1x PBS with 0.1% Triton X-100 for 15 min at RT, incubated in the blocking solution for 15 min at RT, incubated for 1 hour with primary antibodies at RT, washed three times for 10 min with the blocking solution, incubated for 1 hour with secondary antibodies at RT, and washed three times for 10 min in the blocking solution. GV oocytes, eggs, and two-cell embryos were incubated for 30 min with Antifade Mounting Medium with DAPI (Vector Laboratories, cat# H-1200) diluted 1:1 in the blocking solution and placed into 3 μl drops of the blocking solution covered with paraffin oil in a glass-bottom tissue culture dish (fluoroDish, cat# FD35–100). Oocytes at prometaphase I were mounted on microscope slides with Antifade Mounting Medium with DAPI. The following primary antibodies were used: rabbit anti-mouse SP100 antibody (Figures 3A, 3B, and S3, 1:100, custom serum raised by LabCorp), rabbit anti-human SP140 antibody (Figure S6A, 1:100, Invitrogen, cat# PA5-110417), and rabbit anti-Histone H3K9me3 antibody (Figure 2F, 1:100, Abcam, cat# ab8898). Secondary antibodies used were Alexa Fluor 488–conjugated donkey anti-rabbit (1:500, Invitrogen, cat# A21206), and Alexa Fluor 568–conjugated donkey anti-rabbit (1:500, Invitrogen, cat# A10042).

### Immunostaining and Oligopaint FISH of mouse oocytes and embryos

Immunostaining combined with FISH for mouse oocytes and two-cell embryos followed our recently developed protocol.^64^ Briefly, prometaphase I oocytes and two-cell embryos were immunostained with rabbit anti-Histone H3K9me3 antibody (Figure 2F, 1:100, Abcam, cat# ab8898) and Alexa Fluor 568–conjugated donkey anti-rabbit (1:500, Invitrogen, cat# A10042) secondary antibody. Two-cell embryos were immunostained with mouse anti-mouse SP110 (Figure 3C, 1:33, Santa Cruz, cat# sc-376345) or rabbit anti-mouse SP100 antibody (Figure S3, 1:100, custom serum raised by LabCorp) as a primary antibody and Alexa Fluor 488–conjugated donkey anti-rabbit (1:500, Invitrogen, cat# A21206) or Alexa Fluor 488–conjugated donkey anti-mouse (1:500, Invitrogen, cat# A21202) as a secondary antibody. Embryos were incubated overnight at 4°C in the primary antibody for the SP110 staining or 1 hour at RT for the SP100 staining. The blocking solution used for this immunostaining was (0.3% BSA and 0.01% Tween-20 in 1x PBS). Subsequent FISH used 100 pmol of the HEX probe. To improve the retention rate of embryos on the microscope slide, we made the following modification to the original protocol.^64^ Prior to attaching embryos to the slide, embryos were washed in 1x PBS in ultra-low attachment microplates (Corning Costar, cat# 07-200-601) to prevent adhesion to the dish. Embryos were then transferred to a 1x PBS drop on Fisherbrand ColorFrost Plus microscope slides (Fisher Scientific, cat# 12-550-16A), instead of poly-L-lysine treated slides,^64^ allowing them to attach. After PFA fixation,^64^ slides were washed three times for 5 min with 10 ml of 1x PBS while placing the slides horizontally on the bench (instead of inserting them vertically into cuplin jar), which minimizes embryo detachment from the slide. The remaining FISH procedure followed the original protocol.^64^

### Immunostaining and FISH on E6.5 embryos

Following natural mating, embryos were dissected at E6.5 as described above. After dissection, the ectoplacental cones were removed using 27G needles, and embryos were washed in three drops of 1x PBS to avoid contamination of maternal tissues. Embryonic cells were then spread on microscopy slides in the fixative solution (4% PFA and 0.01% Tween-20 in 1x PBS). Cells were mechanically dissociated by pipetting up and down using a mouth-operated pipette. The PFA solution was allowed to dry, and the slides were frozen at −20°C. Each embryo was spread within a ring made with rubber cement (Elmer’s, cat# E904) on the slide using a 1 ml syringe with a 27G needle.

For immunostaining, slides were thawed and briefly washed three times in 1x PBS and incubated in the blocking solution (10% BSA, 0.01% Tween-20 in 1x PBS) for 1 hour at RT in a humidified chamber. The slides were then incubated with a rabbit anti-human 53BP1 antibody (Figure 4A, 1:2000, Invitrogen, cat# PA1-16565) for 1 hour at RT, washed three times with 1x PBS for 10 min, incubated with Alexa Fluor 647–conjugated donkey anti-rabbit (1:500, Invitrogen, cat# A32795) in the blocking solution (2% BSA, 0.01% Tween-20 in 1x PBS), and washed three times with 1x PBS for 10 min. Rubber cement rings were removed using a fine tweezer, and slides were mounted with Antifade Mounting Medium with DAPI. After imaging the immunostained slides, coverslips were removed by immersion in 1x PBS to proceed to the Oligopaint FISH process. Slides were washed for 10 min in 1x PBS, dehydrated in ethanol series (85%, 90%, 100%), air-dried, and aged at RT for one day. Slides were fixed with 1% paraformaldehyde in 1x PBS for 10 min at RT, washed for 5 min with 1x PBS at RT, dehydrated in ethanol series, air dried, and the primary Oligopaint mix^64^ with 100 pmol of the HEX probe was mounted to the slides and sealed with rubber cement. Subsequent steps were performed as described in Clark et al.^64^

### Plasmid construction and cRNA synthesis

Plasmids were synthesized by Twist Bioscience using the plasmid In Vitro Transcription (pIVT) vector^65^ with N-terminal EGFP for SP100-rs, SP100, SP110, and SP140 constructs and pIVT with N-terminal mCherry for the SP110 construct. Plasmid DNA linearized by NdeI (New England Biolabs, cat# R0111S) were used as a template to synthesize cRNAs with the T7 mMessage mMachine Kit (Ambion, cat# AM1340), followed by purification using the MEGAclear Kit (ThermoFisher, cat# AM1908). cRNAs were diluted in nuclease-free water to adjust their concentrations for microinjection (see below).

### Zygote microinjection and immunostaining

Zygotes were microinjected with cRNA in M2 media using a piezo-driven micromanipulator, piezoXpert (Eppendorf). During microinjection, zygotes with abnormal pronuclei numbers and unfertilized eggs were discarded. Following the microinjection, zygotes were cultured to different stages at 37°C in a humidified atmosphere of 5% CO_2_ in air. cRNAs used in this study were *Egfp-Sp100-rs* (*M. m. domesticus* SP100-rs with EGFP at the N- terminus) at 1743 ng/μl, *Egfp-Sp100* (*M. m. domesticus* SP100 (isoform 1) with EGFP at the N - terminus) at 62.5, 90, 125, 250, and 1767 ng/μl, *Egfp-Sp110* (*M. m. domesticus* SP110 with EGFP at the N-terminus) at 125 ng/μl, *mCherry-*Sp110 (*M. m. domesticus* SP110 with mCherry at the N-terminus) at 125 and 250 ng/μl, and *Egfp-Sp140* (*M. m. domesticus* SP140 with EGFP at the N-terminus) at 125 and 250 ng/μl. cRNAs used in co-injections were *Egfp-Sp100* at 90 ng/μl and *mCherry-Sp110* at 125 ng/μl or *Egfp-Sp100* at 125 ng/μl and *mCherry-Sp110* at 250 ng/μl. After culturing, embryos were imaged at E1.5 or E4.5 to examine protein expression levels. Subsequently, embryos were fixed in freshly prepared 2% paraformaldehyde in 1x PBS with 0.1% Triton X-100 for 20 min at RT, washed in blocking solution (0.3% BSA and 0.01% Tween-20 in 1x PBS) for 10 min at RT, permeabilized in 1x PBS with 0.1% Triton X-100 for 15 min at RT, incubated in the blocking solution (5% BSA and 0.01% Tween-20 in 1x PBS) for 1 hour at RT, incubated 1 hour with rabbit anti-human 53BP1 antibody (Figures 4B and 4C, 1:2000, Invitrogen, cat# PA1-16565) at RT, washed three times for 10 min with the blocking solution (5% BSA and 0.01% Tween-20 in 1x PBS), incubated 1 hour with Alexa Fluor 647–conjugated donkey anti-rabbit (1:500, Invitrogen, cat# A32795) antibody at RT, washed three times for 10 min in the blocking solution (0.3% BSA and 0.01% Tween-20 in 1x PBS), incubated 30 min with Antifade Mounting Medium with DAPI (Vector Laboratories, cat# H-1200) diluted 1:1 in the blocking solution, and placed into 3 μl drops of the blocking solution covered with paraffin oil in a glass-bottom tissue culture dish (fluoroDish, cat# FD35–100) to proceed to confocal microscopy (see below).

### Microscopy and image analysis

Fixed oocytes and embryonic cells were imaged with a microscope (Eclipse Ti; Nikon) equipped with 20x / 0.75 NA objective and 100x / 1.40 NA oil-immersion objective lens, CSU-W1 spinning disk confocal scanner (Yokogawa), ORCA Fusion Digital CMOS camera (Hamamatsu Photonics), and 405, 488, 561 and 640 nm laser lines controlled by the NIS-Elements imaging software (Nikon). Confocal images were acquired as Z-stacks at 1 or 3 µm intervals. Fixed oocytes and embryos were imaged with a microscope (Eclipse Ti2-E; Nikon) equipped with the 20x / 0.75 NA objective and 60x / 1.40 NA oil-immersion objective, CSU-W1 spinning disk confocal scanner (Yokogawa), ORCA Fusion Digital CMOS camera (Hamamatsu Photonics), and 405, 488, 561 and 640 nm laser lines controlled by the NIS-Elements imaging software (Nikon). Confocal images were collected as Z-stacks at 0.5, 1, or 3 µm intervals. Fiji/ImageJ (NIH) was used to analyze all the images. In general, optical slices containing nuclei or cells were added to produce a sum intensity Z-projection for pixel intensity quantifications. For quantifications of foci, optical slices were added to produce a sum (Figure 4A) or maximum intensity Z-projection (Figures 4B and 4C).

To quantify H3K9me3 signal intensities at the HEX locus (Figure 2F), regions of interest (ROIs) were drawn to encompass the H3K9me3 signal on the chromosome 1 arm. In general, H3K9me3 signals are enriched at the centromeric region of each chromosome, and the only H3K9me3 focus outside the centromere corresponds to the HEX locus. The signal intensity was integrated over each ROI after subtracting background signals, obtained from the average of three chromosome regions near the HEX locus.

To quantify SP100 nucleoplasmic signal intensities in meiosis I GV oocytes and two-cell embryos (Figure 3A), ROIs were drawn encompassing the SP100 nuclear signal based on the DAPI staining. The signal intensity was integrated over each ROI after subtracting background signals, obtained from the background outside the cell. To quantify SP100 and SP140 total signal intensities in meiosis II eggs and two-cell embryos (Figures 3B and S6A), ROIs were drawn encompassing the entire cell. The signal intensity was integrated over each ROI after subtracting background signals, obtained from three background areas outside the cell. To quantify nucleoplasmic signal intensities in two-cell embryos (Figure 3B), masking images were created using DAPI staining images to specifically measure signal intensities in the nucleus. The signal intensity was integrated over each slice after subtracting background signals, obtained from three background areas outside of the cell. To quantify nucleoplasmic signal intensities (SP100 and SP110) in IF-FISH images (Figures 3C and S3), masking images were created using DAPI staining images to specifically measure signal intensities in the nucleus. Signal intensity was averaged over each slice after subtracting cytoplasmic background signals. Nucleoplasmic signals of EGFP-SP100 (Figure 4C) were quantified by the same method.

To quantify 53BP1 foci number per cell (Figure 4A), we developed a custom ImageJ/Fiji macro (Supplementary Text) based on the strategy developed by the Duke Light Microscopy Core (https://microscopy.duke.edu/guides/count-nuclear-foci-ImageJ). We manually counted 53BP1 foci number in Figures 4B and 4C, using maximum-intensity projected images. Foci with ≥ 1 μm diameter were counted. We analyzed each slice when nuclei from different cells overlap.

### Embryo transfer

Embryo transfers were performed by the NHLBI Transgenic Core at NIH. In the first embryo transfer experiment (Figure 1G), +/+ zygotes were collected from +/+ females mated with +/+ males and cultured for one day to reach the two-cell stage. Two-cell embryos were then transferred to pseudo-pregnant H/+ *dom* females (mated with vasectomized males prior to the transfer). As a control, embryos were also transferred to pseudo-pregnant +/+ females. Embryos were dissected from these females at E9.5 (two +/+ and three H/+ females). The number of live embryos and resorbing sites were scored to quantify embryo survival rates in the H/+ and +/+ uterus. A replica of this experiment was performed, and the embryos were dissected at E13.5 and E17.5 (three +/+ and three H/+ females for each timepoint).

In a second embryo transfer experiment (Figure 1H), a mixture of +/+ and H/+ two-cell embryos obtained by crossing H/+ *dom* females with +/+ *dom* males were transferred to eight +/+ pseudo-pregnant *dom* females (CD-1 background). Embryos were dissected at E13.5 and genotyped by qPCR as previously described. In a replica of this experiment, two-cell embryos were implanted into five pseudo-pregnant females and dissected and genotyped at E14.5.

### Gene expression profiling by RT-qPCR

Quantification of expression levels of *Sp100*, *Sp110*, *Sp140*, and *Sp100-rs* genes were performed using liver and oocytes of *M. m. domesticus* and *M. m. musculus.* Dissected tissues were immediately frozen in liquid nitrogen and stored at −70°C. RNA from oocytes was extracted using the TRIzol Reagent (Invitrogen, cat# 15596026) and from liver using the total RNA extraction kit (Norgen, cat#17200). RNA integrities were examined on 1% agarose gels. RNA purities were examined using the NanoPhotometer NP80 (Implen, cat# NC2831907). cDNA for each sample was synthesized using the SuperScript™ III First-Strand Synthesis System (Invitrogen, cat# 18080051) using 0.8 μg per sample of total RNA, following the manufacturer’s instructions. Working solutions of the obtained cDNAs were produced by diluting them 1:5 in RNase-DNase free water.

Primers were designed using a modified version of Primer3^61^ implemented in Geneious 10.2.6. Primer sets for *Sp100*, *Sp110*, *Sp140*, and *Sp100-rs* genes (Table S1) were designed spanning an exon-exon junction of each gene. Primer specificities were confirmed using the Blast algorithm for short sequences.^61,66^ *Sp100-rs* gene is a chimeric gene with the first three exons of *Sp100* fused with *Csprs* gene. To specifically amplify *Sp100-rs*, we designed a primer set spanning the junction of the 3^rd^ exon of *Sp100* and the exon of *Csprs*, which avoids amplifying *Sp100* (Figure S5C). Quantitative PCR was performed on QuantStudio 3 (Applied Biosystems, cat# A28567), except for *Sp140* in *dom* oocytes samples that were performed on LightCycler 96. Reactions were assembled in a final volume of 10 µl with 1 μl of cDNA, 5 μl of iTaq Universal SYBR Green Supermix (Bio-Rad, cat# 1725121), and 0.5 μl of each 10 µM primer. Cycle conditions were 95°C for 10 min; 40 cycles of 95°C for 5 s and 60°C for 20 s. The specificity of the PCR products was confirmed by dissociation curve analysis. Target and reference genes were analyzed simultaneously in triplicates for independent samples. DNA contamination was examined using controls without reverse transcriptase (NRTC) and without template cDNA (NTC). A five-fold serial dilution of the cDNAs was used to calculate the primer sets amplification efficiency (E). An efficiency range of 90% to 110% was used to select primer sets for further analysis. Cq values were corrected using the gene’s efficiencies with the formula CqE = Cq*(log(1+E)/log(2)).^67^ The normalized relative expression quantity (NREQ) was determined by the 2^−ΔΔ*C*^_T_ method.^68^ Expression levels of target genes were normalized using the geometric mean of two housekeeping reference genes with subsequent normalization to the average expression of the +/+ group. To select the reference genes, we used the RefFinder^69^ software. From five tested genes (*Atp5b*, *Gapdh*, *Hprt1*, *Rplpo*, and *Ubc*), we selected *Atp5b* and *Rplpo* for *dom* liver, *Atp5b* and *Gapdh* for *mus* liver, *Atp5b* and *Hprt1* for *dom* and *mus* oocytes.

## Quantification and statistical analysis

Data points were pooled from more than two independent experiments in most experiments, and the exact number of independent experiments for each experimental group is listed in Table S2. Data was analyzed using Microsoft Excel and GraphPad Prism 10. Bar graphs, scatter plots, and violin plots were created with GraphPad Prism 10. Binomial test (two-sided) for deviations from the expected 50:50 (Figure 1D and Table 1) or the biasing cross ratio (Figure 1H), Chi-square test (two-tailed) (Figure 1G and Table 2), Mann-Whitney test (two-sided) (Figures 1E, 1F, 2B–2E, 3A, 3B nucleoplasmic, 3C, 4A, S3, and S6A), and Welch’s t test (two-sided) (Figures 2F, 3B total, and 4C) were used for statistical analyses. Actual *P* values are shown in each figure or figure legend.

## Supplementary Text

Custom ImageJ/Fiji macro used to quantify 53BP1 foci in Figure 4A:

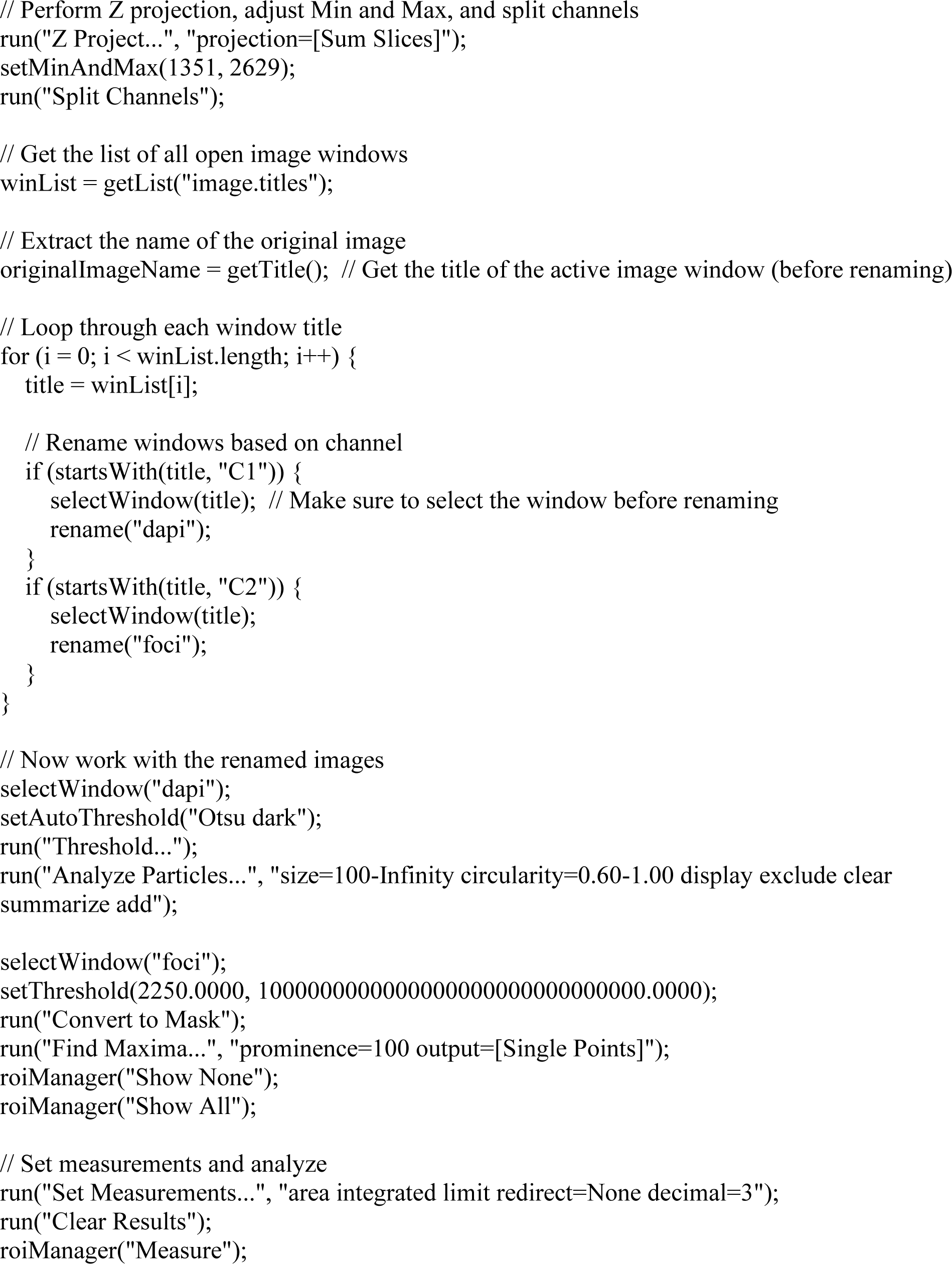
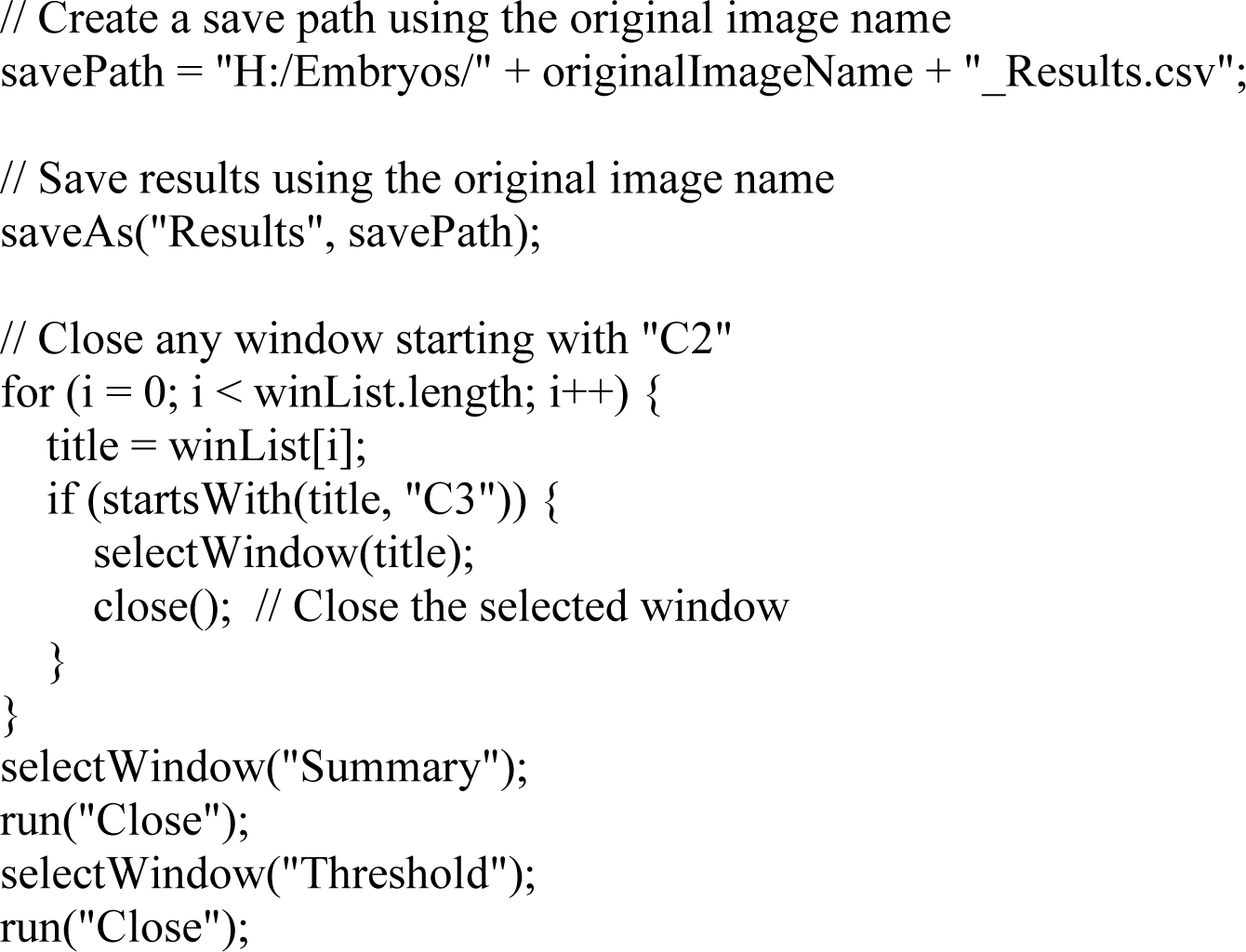

---

**Figure S1.**
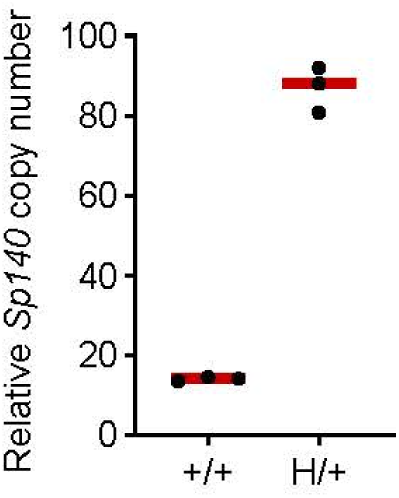
Relative copy number of *Sp140* in *Mus musculus domesticus* estimated by qPCR, related to. Figure 2. DNA samples extracted from tail snips were used to examine the copy number increase of *Sp140* gene in H/+ mice compared to +/+. See Method details “DNA isolation and genotyping by qPCR”.

**Figure S2.**
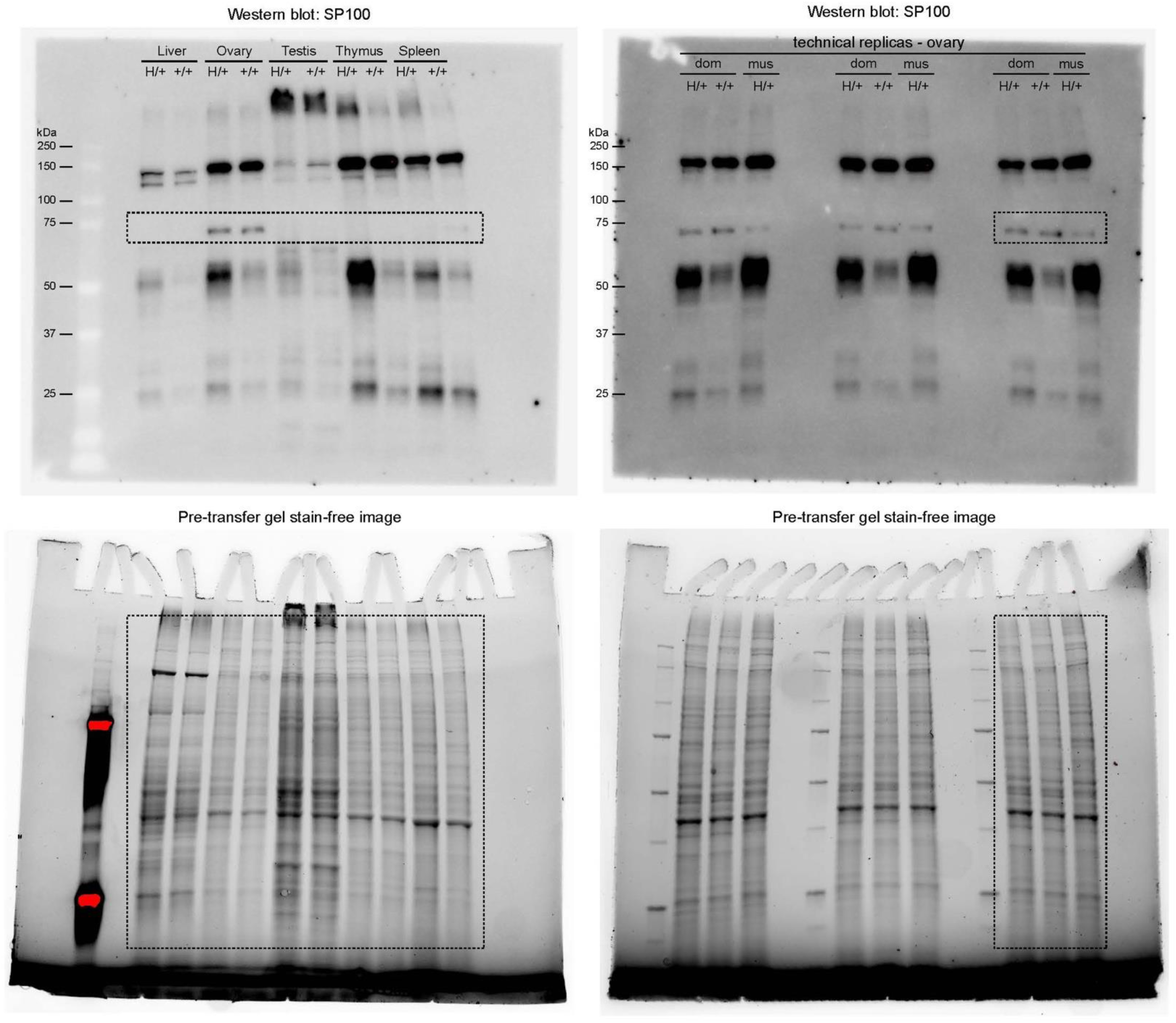
Uncropped images of western blots, related to. Figure 2. Entire gels and membranes are imaged. Boxes in the images indicate the cropped area shown in Figure 2G.

**Figure S3.**
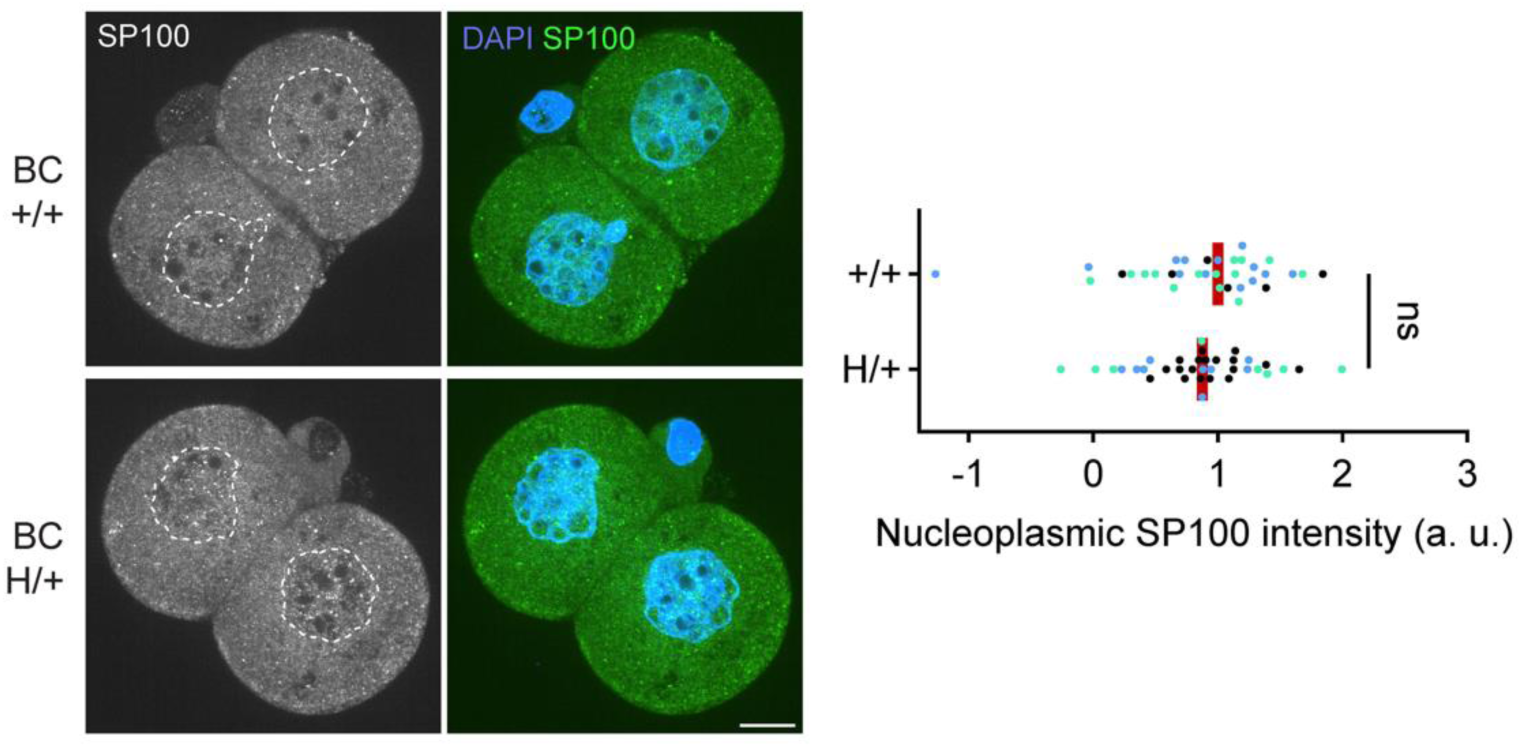
SP100 toxin is equally deposited to +/+ and H/+ embryos, related to. Figure 3. Two-cell embryos from BC were fixed, stained for SP100 and genotyped by FISH with the HEX probe. SP100 signal intensities in the nucleus were quantified (n = 33, and 35 for +/+ and H/+, respectively). Mann-Whitney test (two-sided) was used for statistical analysis; ns = 0.5338; red line, median; dots, individual cells. This experiment was repeated independently three times, indicated by dots with distinct colors. Images are optical slices; scale bar, 10 *μ*m.

**Figure S4.**
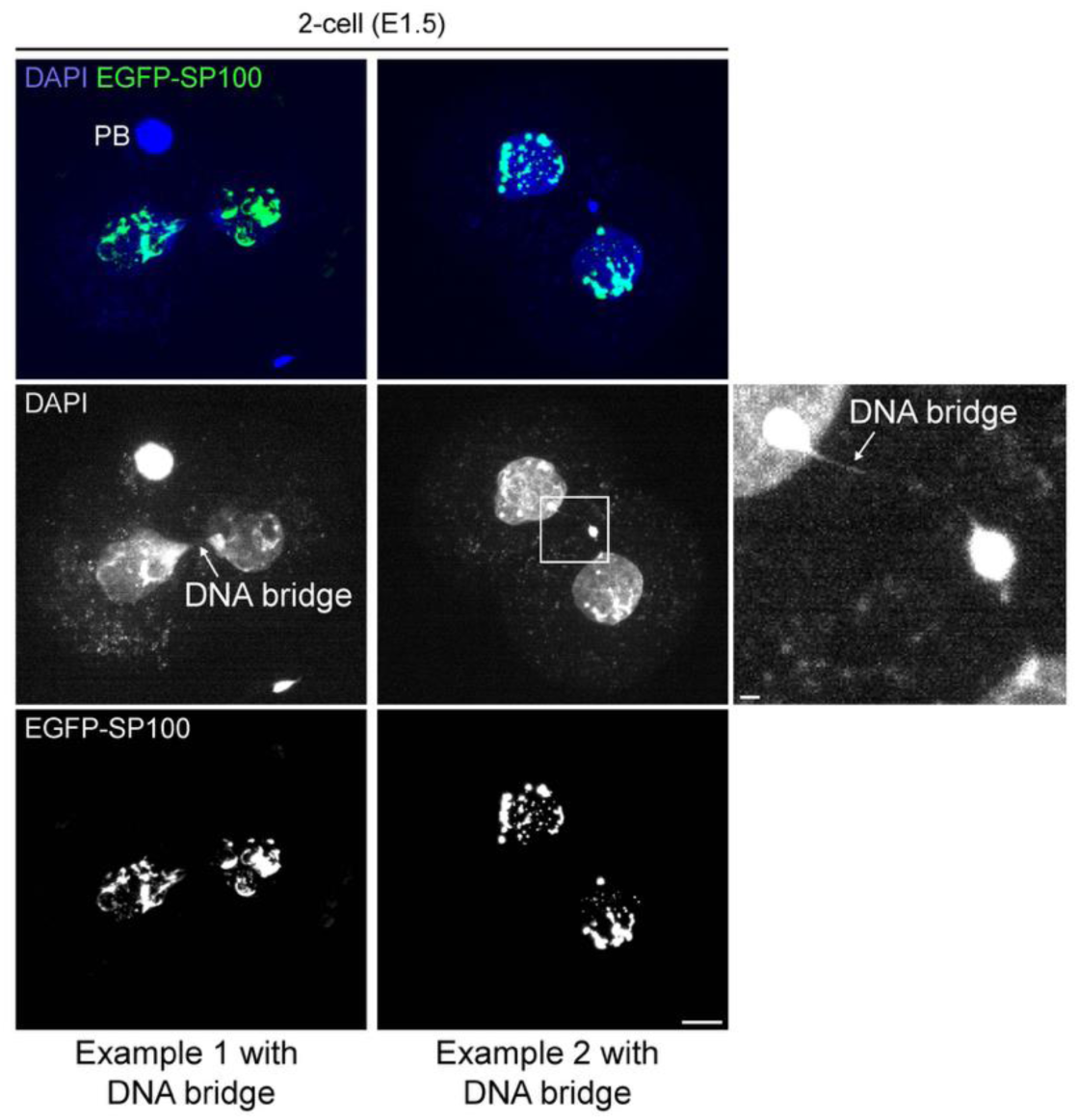
DNA bridges induced upon SP100 overexpression in embryos, related to. Figure 4. Embryos overexpressing EGFP-SP100 were fixed at the two-cell stage; Images are maximum intensity Z-projections; E, embryonic day; PB, polar body; scale bar, 10 *μ*m; scale bar of enlarged inset, 1 *μ*m

**Figure S5.**
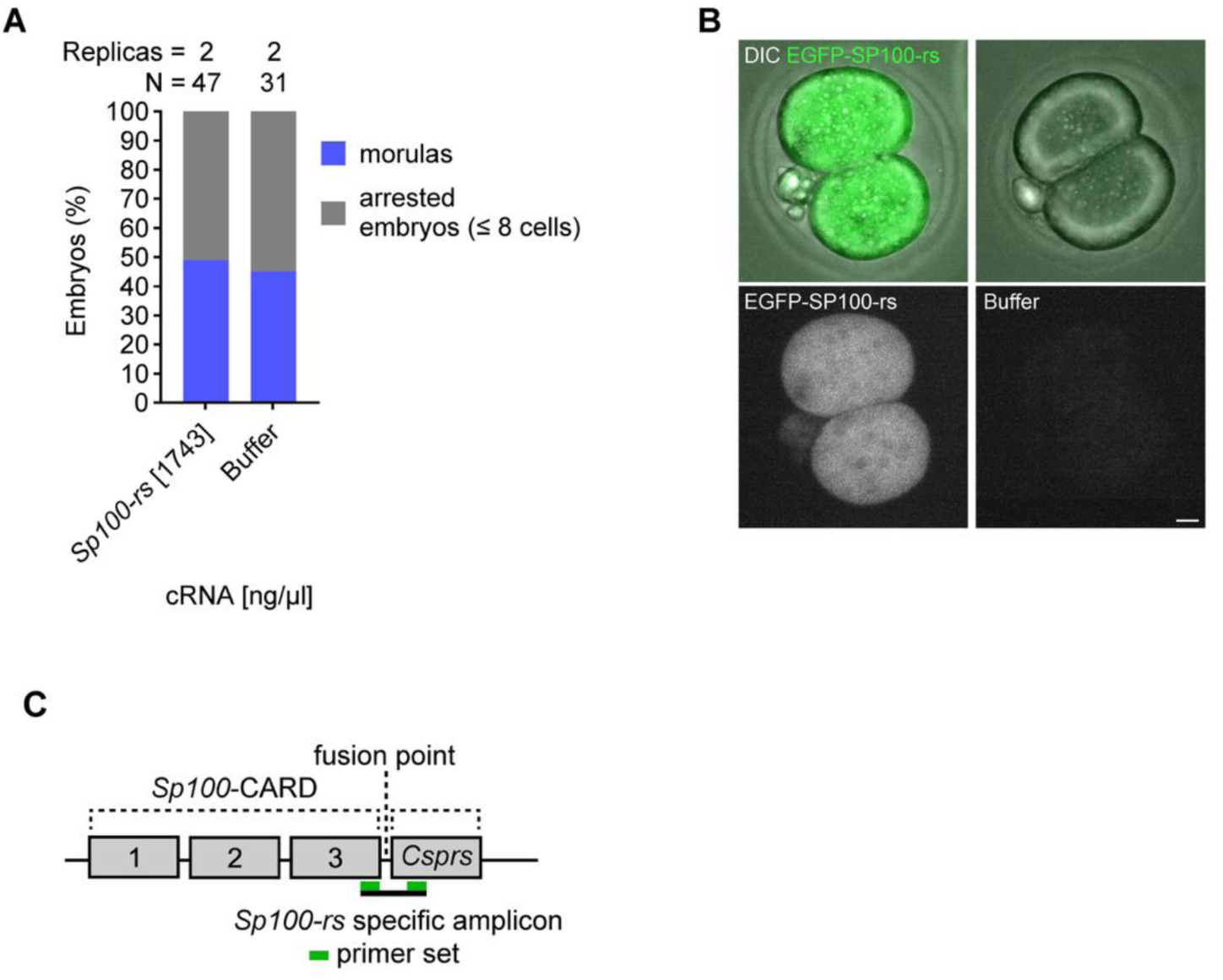
No significant impact on early embryogenesis upon *Sp100-rs* overexpression, related to. Figure 4**. (A)** Control zygotes were microinjected with *Egfp-Sp100-rs* cRNA (1743 ng/μl) and cultured for three days. Embryos were categorized as normal morula (blue) or arrested embryos (grey); sample size is shown above each bar; data from two independent experiments were pooled. **(B)** Control zygotes were microinjected with *Egfp-Sp100-rs* cRNA (1743 ng/μl) or buffer and cultured for one day (the two-cell stage) to examine SP100-rs localization. Ectopically expressed EGFP-SP100-rs uniformly localized in the cell and did not show nuclear enrichment in contrast to SP100 (Figure 4C). Images are maximum intensity Z-projections; scale bar, 10 *μ*m. **(C)** Primer designs to specifically amplify *Sp100-rs*.

**Figure S6.**
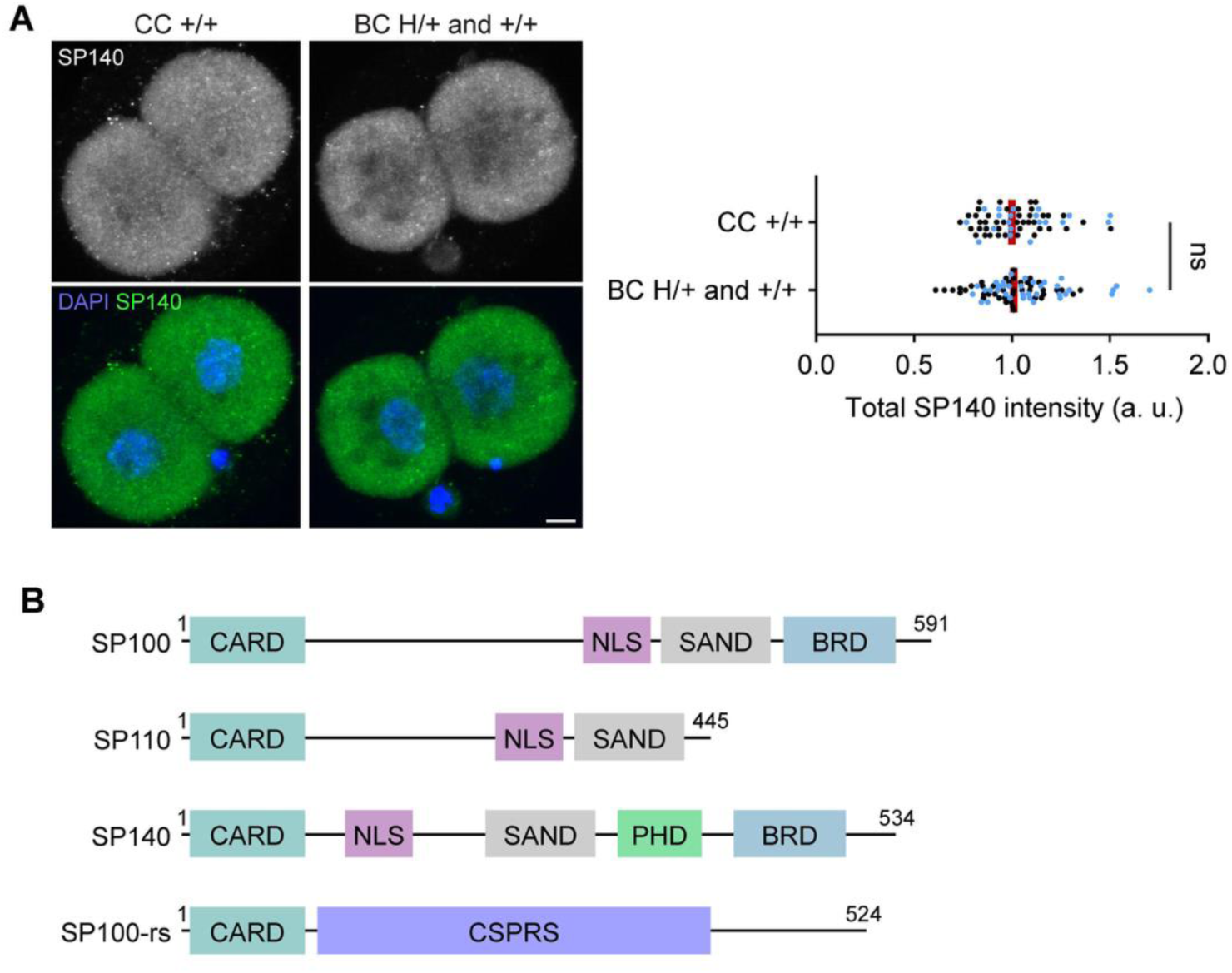
SP140 levels are indistinguishable in two-cell embryos from the control and biasing crosses, related to. Figure 4**. (A)** Embryos from the control cross (CC) and the biasing cross (BC) were fixed at the early two-cell stage and stained for SP140. The signal intensities in the cell were quantified (n = 62 and 78 for +/+ CC and H/+ and +/+ BC, respectively). Mann-Whitney test (two-sided) was used for statistical analysis; ns = 0.7372; red line, median; dots, individual cells from two-cell embryos. Data points were pooled from two independent experiments, indicated by dots with distinct colors. Images are maximum intensity Z-projections; scale bar, 10 *μ*m. **(B)** Domain organizations of the Speckled Protein (SP) family: CARD (Caspase Activation and Recruitment Domain) also known as HSR domain; NLS (Nuclear Localization Signal); SAND (named after its host proteins: SP100, Aire, NucP41/P75, and DEAF); PHD (Plant Homeodomain); BRD (Bromodomain); CSPRS (Component of *Sp100-rs*). SP100-rs does not have NLS, consistent with its uniform distribution in the cell when overexpressed (Figure S5B).

**Table S1.**
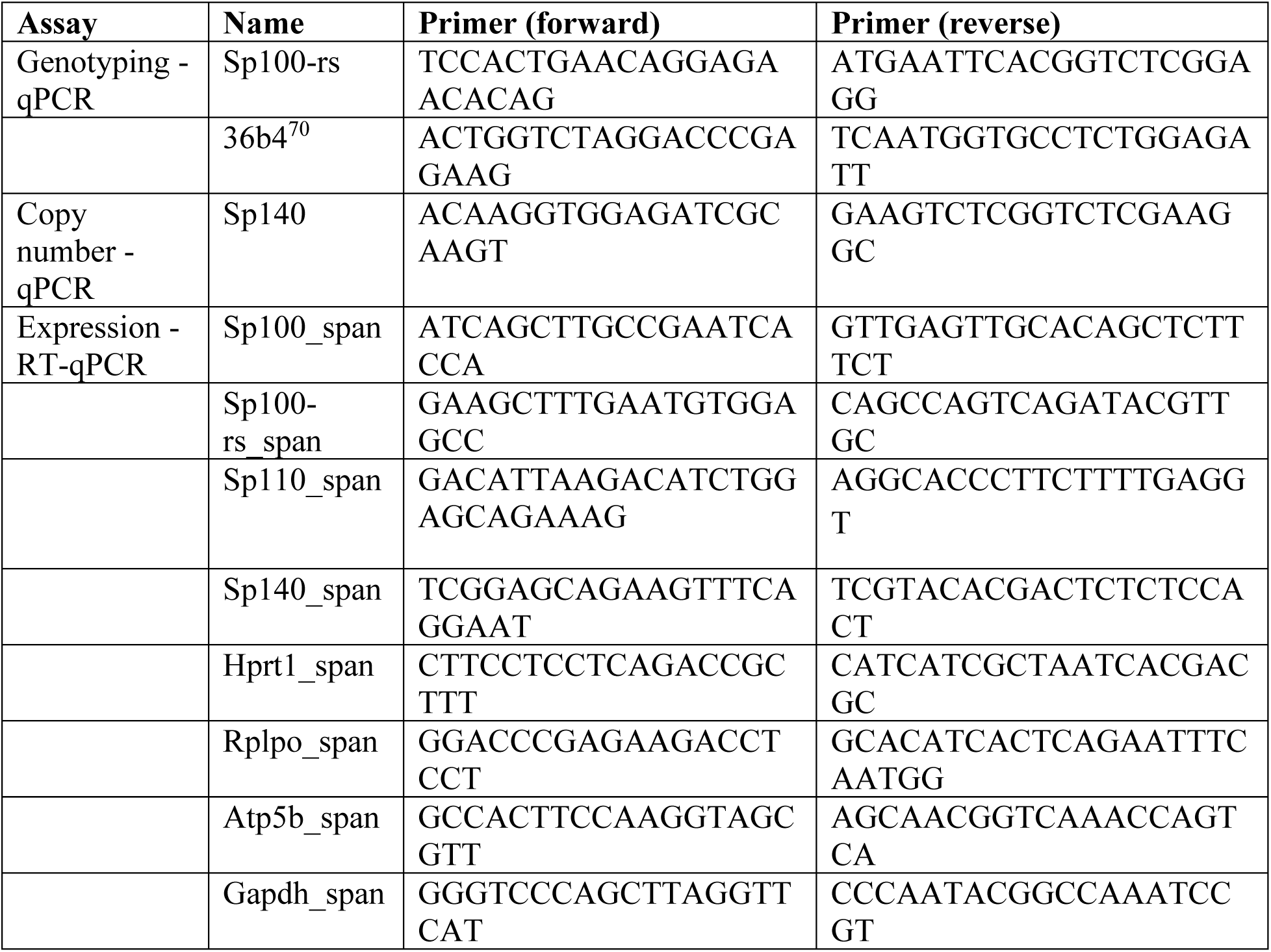
List of primer sets used in this study.

**Table S2.**
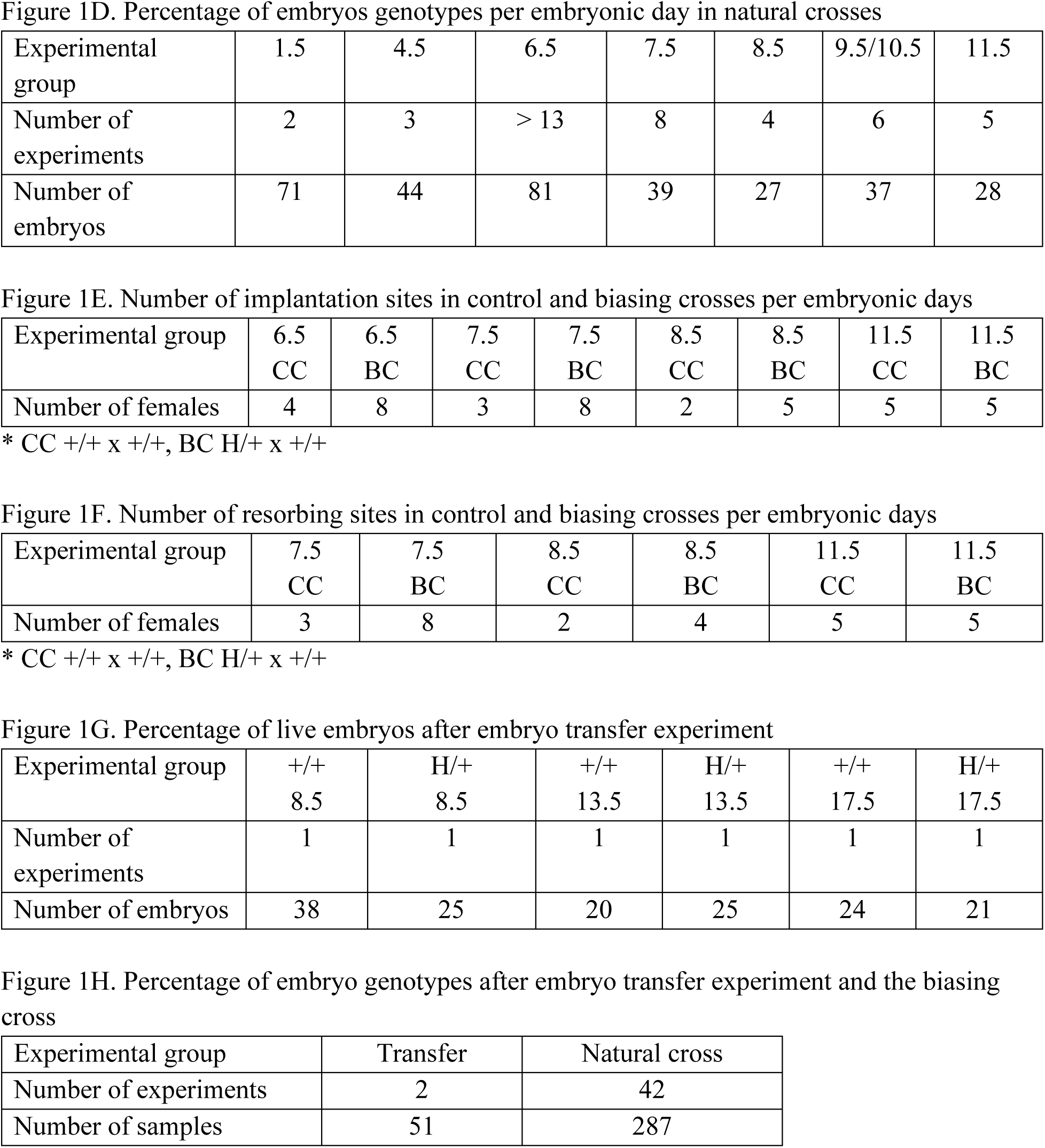
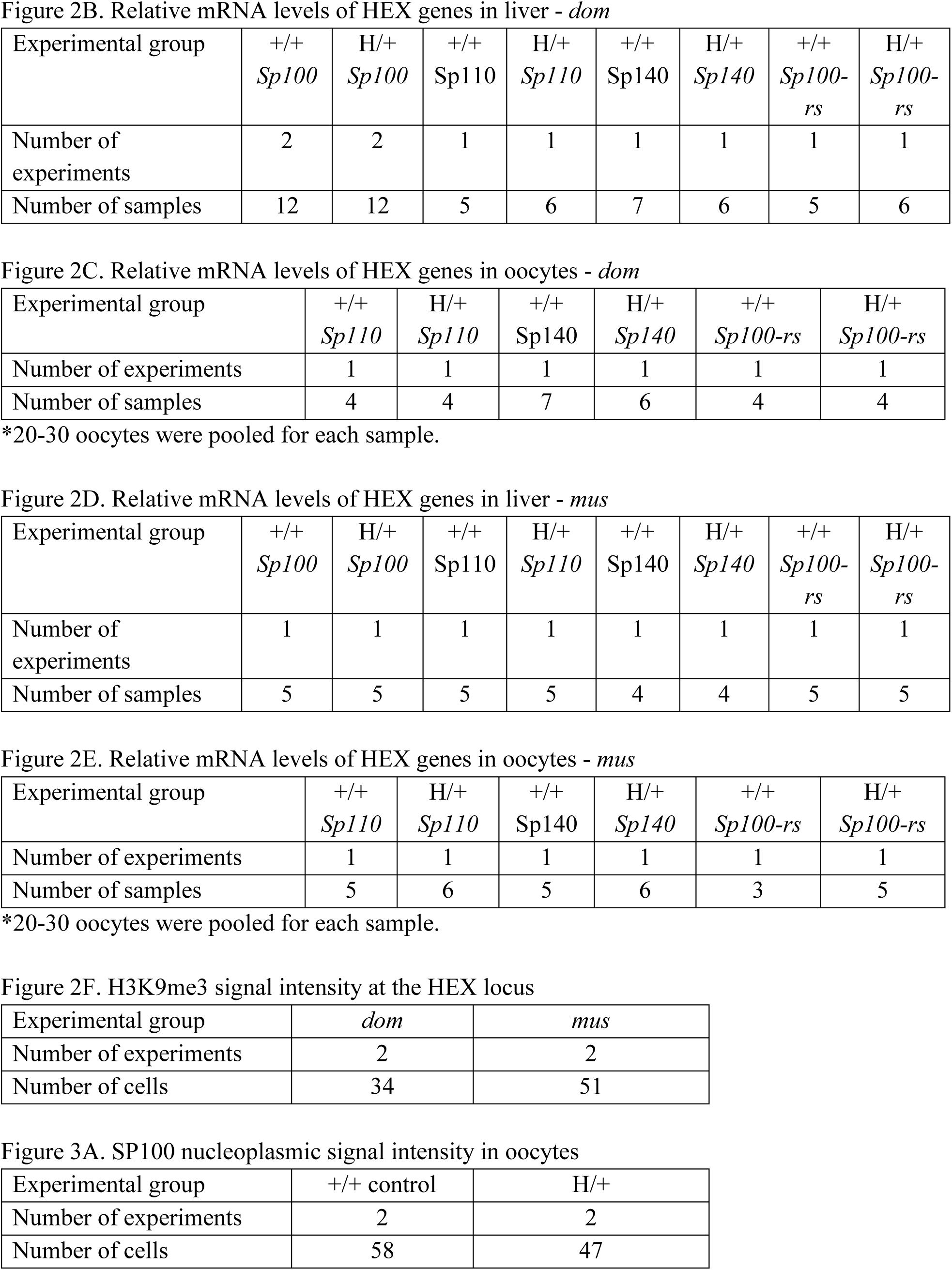
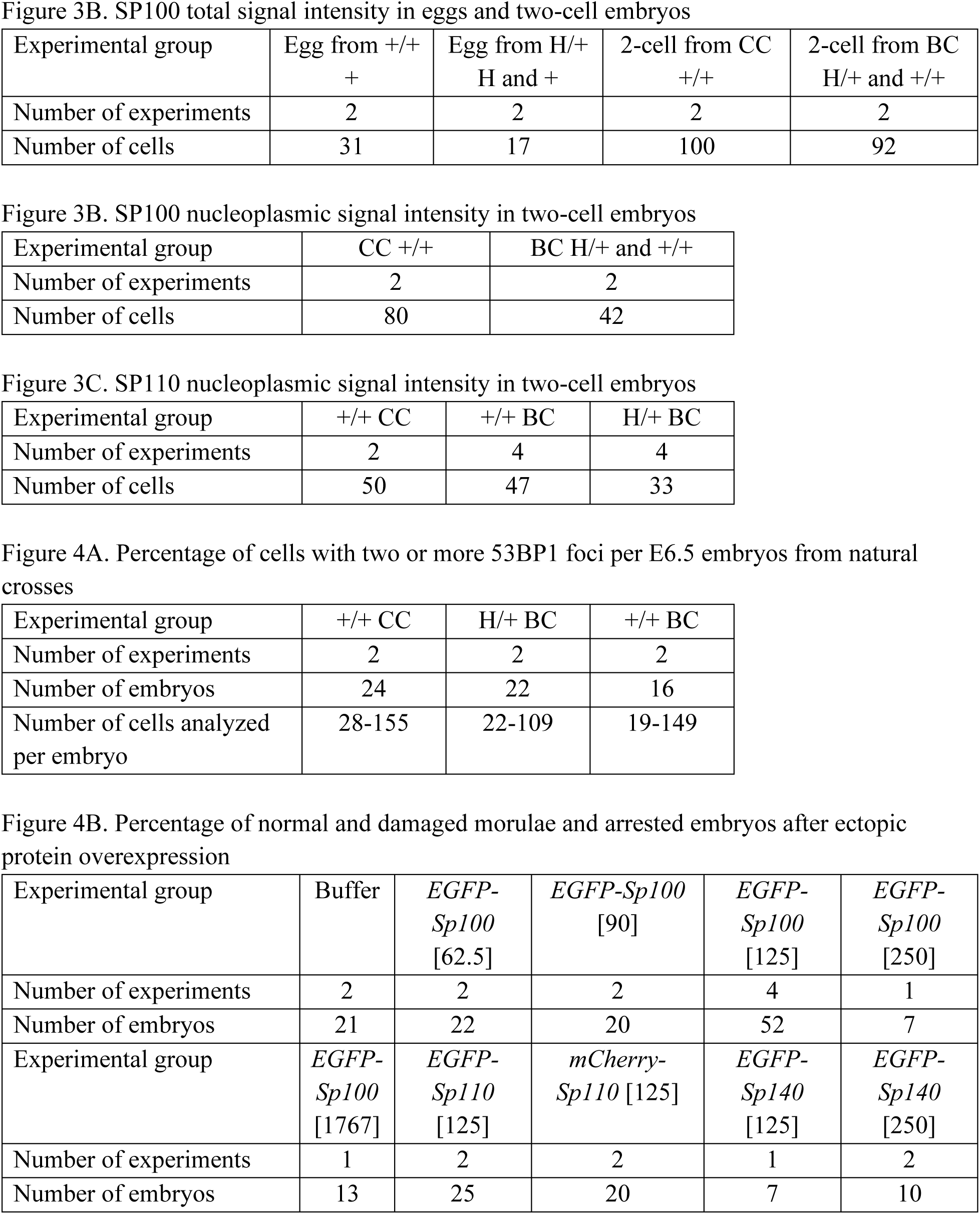
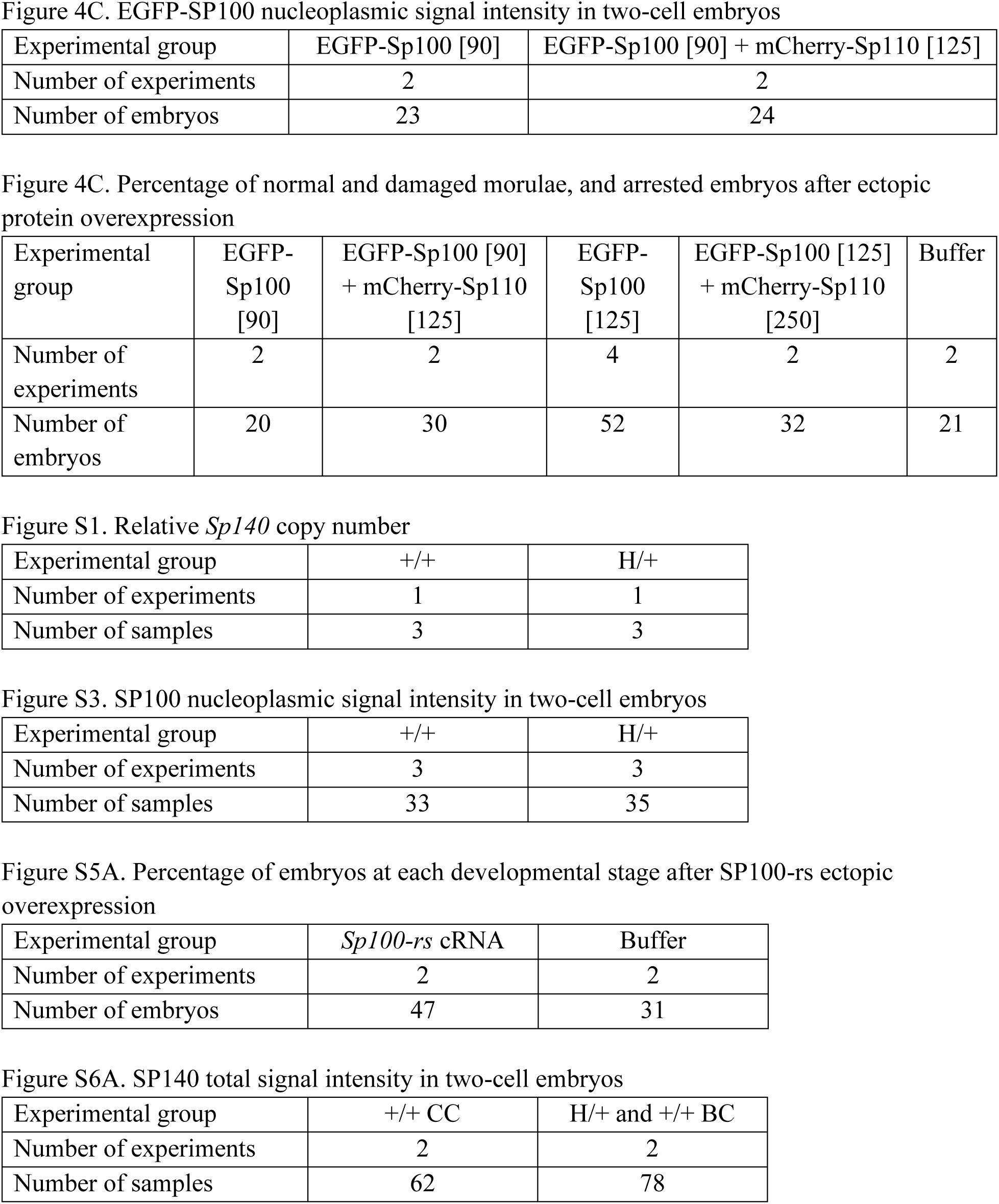
List of the total numbers of events analyzed and the numbers of repeats of each experiment.

**Figure 1D. Percentage of embryos genotypes per embryonic day in natural crosses**
| Experimental group | 1.5 | 4.5 | 6.5 | 7.5 | 8.5 | 9.5/10.5 | 11.5 |
| --- | --- | --- | --- | --- | --- | --- | --- |
| Number of experiments | 2 | 3 | > 13 | 8 | 4 | 6 | 5 |
| Number of embryos | 71 | 44 | 81 | 39 | 27 | 37 | 28 |

**Figure 1E. Number of implantation sites in control and biasing crosses per embryonic days**
| Experimental group | 6.5<br>CC | 6.5<br>BC | 7.5<br>CC | 7.5<br>BC | 8.5<br>CC | 8.5<br>BC | 11.5<br>CC | 11.5<br>BC |
| --- | --- | --- | --- | --- | --- | --- | --- | --- |
| Number of females | 4 | 8 | 3 | 8 | 2 | 5 | 5 | 5 |

\* CC +/+ x +/+, BC H/+ x +/+

**Figure 1F. Number of resorbing sites in control and biasing crosses per embryonic days**
| Experimental group | 7.5<br>CC | 7.5<br>BC | 8.5<br>CC | 8.5<br>BC | 11.5<br>CC | 11.5<br>BC |
| --- | --- | --- | --- | --- | --- | --- |
| Number of females | 3 | 8 | 2 | 4 | 5 | 5 |

\* CC +/+ x +/+, BC H/+ x +/+

**Figure 1G. Percentage of live embryos after embryo transfer experiment**
| Experimental group | +/+<br>8.5 | H/+<br>8.5 | +/+<br>13.5 | H/+<br>13.5 | +/+<br>17.5 | H/+<br>17.5 |
| --- | --- | --- | --- | --- | --- | --- |
| Number of experiments | 1 | 1 | 1 | 1 | 1 | 1 |
| Number of embryos | 38 | 25 | 20 | 25 | 24 | 21 |

**Figure 1H. Percentage of embryo genotypes after embryo transfer experiment and the biasing cross**
| Experimental group | Transfer | Natural cross |
| --- | --- | --- |
| Number of experiments | 2 | 42 |
| Number of samples | 51 | 287 |

| Experimental group | +/+ *Sp100* | H/+ *Sp100* | +/+ *Sp110* | H/+ *Sp110* | +/+ *Sp140* | H/+ *Sp140* | +/+ *Sp100-rs* | H/+ *Sp100-rs* |
| --- | --- | --- | --- | --- | --- | --- | --- | --- |
| Number of experiments | 2 | 2 | 1 | 1 | 1 | 1 | 1 | 1 |
| Number of samples | 12 | 12 | 5 | 6 | 7 | 6 | 5 | 6 |

| Experimental group | +/+ *Sp110* | H/+ *Sp110* | +/+ *Sp140* | H/+ *Sp140* | +/+ *Sp100-rs* | H/+ *Sp100-rs* |
| --- | --- | --- | --- | --- | --- | --- |
| Number of experiments | 1 | 1 | 1 | 1 | 1 | 1 |
| Number of samples | 4 | 4 | 7 | 6 | 4 | 4 |

\*20-30 oocytes were pooled for each sample.

| Experimental group | +/+ *Sp100* | H/+ *Sp100* | +/+ *Sp110* | H/+ *Sp110* | +/+ *Sp140* | H/+ *Sp140* | +/+ *Sp100-rs* | H/+ *Sp100-rs* |
| --- | --- | --- | --- | --- | --- | --- | --- | --- |
| Number of experiments | 1 | 1 | 1 | 1 | 1 | 1 | 1 | 1 |
| Number of samples | 5 | 5 | 5 | 5 | 4 | 4 | 5 | 5 |

| Experimental group | +/+ *Sp110* | H/+ *Sp110* | +/+ *Sp140* | H/+ *Sp140* | +/+ *Sp100-rs* | H/+ *Sp100-rs* |
| --- | --- | --- | --- | --- | --- | --- |
| Number of experiments | 1 | 1 | 1 | 1 | 1 | 1 |
| Number of samples | 5 | 6 | 5 | 6 | 3 | 5 |

\*20-30 oocytes were pooled for each sample.

| Experimental group | *dom* | *mus* |
| --- | --- | --- |
| Number of experiments | 2 | 2 |
| Number of cells | 34 | 51 |

| Experimental group | +/+ control | H/+ |
| --- | --- | --- |
| Number of experiments | 2 | 2 |
| Number of cells | 58 | 47 |

| Experimental group | Egg from +/+<br>+ | Egg from H/+<br>H and + | 2-cell from CC<br>+/+ | 2-cell from BC<br>H/+ and +/+ |
| --- | --- | --- | --- | --- |
| Number of experiments | 2 | 2 | 2 | 2 |
| Number of cells | 31 | 17 | 100 | 92 |

| Experimental group | CC +/+ | BC H/+ and +/+ |
| --- | --- | --- |
| Number of experiments | 2 | 2 |
| Number of cells | 80 | 42 |

| Experimental group | +/+ CC | +/+ BC | H/+ BC |
| --- | --- | --- | --- |
| Number of experiments | 2 | 4 | 4 |
| Number of cells | 50 | 47 | 33 |

| Experimental group | +/+ CC | H/+ BC | +/+ BC |
| --- | --- | --- | --- |
| Number of experiments | 2 | 2 | 2 |
| Number of embryos | 24 | 22 | 16 |
| Number of cells analyzed<br>per embryo | 28-155 | 22-109 | 19-149 |

| Experimental group | Buffer | *EGFP-Sp100*<br>[62.5] | *EGFP-Sp100*<br>[90] | *EGFP-Sp100*<br>[125] | *EGFP-Sp100*<br>[250] |
| --- | --- | --- | --- | --- | --- |
| Number of experiments | 2 | 2 | 2 | 4 | 1 |
| Number of embryos | 21 | 22 | 20 | 52 | 7 |
| Experimental group | *EGFP-Sp100*<br>[1767] | *EGFP-Sp110*<br>[125] | *mCherry-Sp110* [125] | *EGFP-Sp140*<br>[125] | *EGFP-Sp140*<br>[250] |
| Number of experiments | 1 | 2 | 2 | 1 | 2 |
| Number of embryos | 13 | 25 | 20 | 7 | 10 |

| Experimental group | EGFP-Sp100 [90] | EGFP-Sp100 [90] + mCherry-Sp110 [125] |
| --- | --- | --- |
| Number of experiments | 2 | 2 |
| Number of embryos | 23 | 24 |

| Experimental group | EGFP-Sp100 [90] | EGFP-Sp100 [90] + mCherry-Sp110 [125] | EGFP-Sp100 [125] | EGFP-Sp100 [125] + mCherry-Sp110 [250] | Buffer |
| --- | --- | --- | --- | --- | --- |
| Number of experiments | 2 | 2 | 4 | 2 | 2 |
| Number of embryos | 20 | 30 | 52 | 32 | 21 |

| Experimental group | +/+ | H/+ |
| --- | --- | --- |
| Number of experiments | 1 | 1 |
| Number of samples | 3 | 3 |

| Experimental group | +/+ | H/+ |
| --- | --- | --- |
| Number of experiments | 3 | 3 |
| Number of samples | 33 | 35 |

| Experimental group | *Sp100-rs* cRNA | Buffer |
| --- | --- | --- |
| Number of experiments | 2 | 2 |
| Number of embryos | 47 | 31 |

| Experimental group | +/+ CC | H/+ and +/+ BC |
| --- | --- | --- |
| Number of experiments | 2 | 2 |
| Number of samples | 62 | 78 |

